# A combinatorial DNA origami platform for biologically replicable, thermostable data storage and molecular authentication

**DOI:** 10.64898/2026.06.08.730396

**Authors:** Ferenc Fördös, Alexander Kloosterman, Alexander Lindberg, Boxuan Shen, Igor Baars, Björn Högberg

## Abstract

DNA origami is becoming an attractive platform for data storage, yet current approaches rely on the limited stability of DNA hybridization, preventing them from fully utilizing the stability and cost-effective copying inherent to classical sequence-based storage. Here we introduce DNA Origami for Combinatorial data Storage (DOCS), where we encode information into the scaffold molecule using a combinatorial enzymatic approach. This enables text encoding that is biologically cloneable, stable at high temperatures, and randomly accessible. We further demonstrate the DOCS platform’s combinatorial power by creating a stochastic molecular authentication system. Finally, we show using simulations that expanding the information capacity of data carriers allows for the storage and recovery of large files up to several hundred kilobytes in size. DOCS provides a robust, scalable strategy for molecular data storage and security that bridges the gap between classical DNA data storage strategies and DNA nanostructure-based methods.

## Main

Inspired by its role as a biological information carrier, DNA has been pursued to be a medium for digital information storage for more than a decade^1^. It has been demonstrated that DNA can provide us with a storage technology with unprecedentedly high information density^2,3^ and extraordinary stability^4,5^ while allowing for data processing operations, such as random access^6^, similarity-based search^7^, data manipulation^8^, and other types of computations on the stored data^9^. As one of the main hurdles for the practical applicability of the sequence-based storage approaches is the prohibitively high costs and slow speed of de novo DNA synthesis^10^, the community has focused on developing alternative, more efficient and scalable ways for synthetizing DNA molecules at demand using enzymatic approaches^11–13^, and chemical arrays^14,15^. Although significant improvements have been achieved in this respect, these technologies are still falling several magnitudes short of the synthesis costs and speeds required for commercial viability on large scales.

Alternative strategies have emerged using predefined sets of DNA molecules to encode information to address this bottleneck^16–19^. One of these uses DNA origami, a versatile method to generate nanostructures with precisely engineerable nanoscale features. DNA origami has become frequently used in physics, biology and medicine among other fields, with applications ranging from creating metamaterials^20^ through investigating receptor signaling^21^ to engineering smart drug carriers^22^. The suitability of DNA origami for storing information has been demonstrated through a number of works that have shown that it can be used in high information density storage platforms^23^ and it allows for data encryption^24^, random access and dynamic data modification via linked memory architectures^25^. Even though these works have shown that the highly engineerable nature of DNA origami structures can be exploited to create a more scalable DNA-based data storage technology, in all of them the information is encoded in the pattern of individual staple oligonucleotides hybridized to the scaffold molecule. This interaction is relatively unstable, preventing one to take full advantage of the key aspects DNA provides for data storage: high degree of stability and low-cost copying using the existing cellular machinery for DNA replication.

Here, we introduce DNA Origami for Combinatorial Storage (DOCS), a technology that encodes information directly into the scaffold sequence via combinatorial enzymatic assembly. In contrast to other approaches, DOCS uses a universal set of staple oligonucleotides to fold distinct, data-encoding scaffold molecules into a library of data carrier origami structures, from which data is read by Exchange-PAINT imaging. As a proof-of-concept, we achieved a functional storage density of 2.4Gb/mm^2^ and successfully encoded and recovered a 67-bit text message with random-access capabilities. Furthermore, we demonstrate how DOCS overcomes the limitations of previous origami-based storage approaches: we successfully recovered DOCS-encoded information after incubation at 80°C for one hour and achieved biological amplification using bacteria. To demonstrate the system’s scalability beyond these proof-of-concept experiments, we used simulations based on experimentally derived error rates to confirm that by utilizing compression (zlib) and expanding the number of imaging channels and signals per data carrier, DOCS can be scaled to store texts ranging from poems to books. Finally, we show that the combinatorial power of the DOCS platform can also be harnessed to create a molecular authentication system. By combining the highly parallel readout and modular construction of DNA origami with the thermostability and biological replicability of sequence-based approaches, DOCS offers a scalable method for both storing digital information and constructing authentication systems using DNA nanostructures.

### Combinatorial data storage in DNA origami using DOCS

DOCS encodes information by combinatorially assembling inserts that are subsequently cloned into the M13 vector. Each insert contains a data payload of five sections, called Data Encoder Fragments (DEFs). A DEF can be any of six possible Data Encoding Sequences (DES; labelled with indices 0-5 from here on out), allowing for the encoding of data in a senary (6-ary) format. Each DES consists of two pairs of Exchange PAINT docking sites that make it uniquely recognizable. In addition, the insert contains an unpaired Data ID sequence. Single stranded information-encoding scaffolds, obtained through phage preparation, can be folded into DNA origami structures called data carriers (DC). Each DC displays their payload as single-stranded DNA, allowing for detection by Exchange-PAINT. The Data ID sequence identifies DPs encoding the same file and can be used for the random access of the information stored (Fig1.a, Fig.S1-2).

**Fig. 1:**
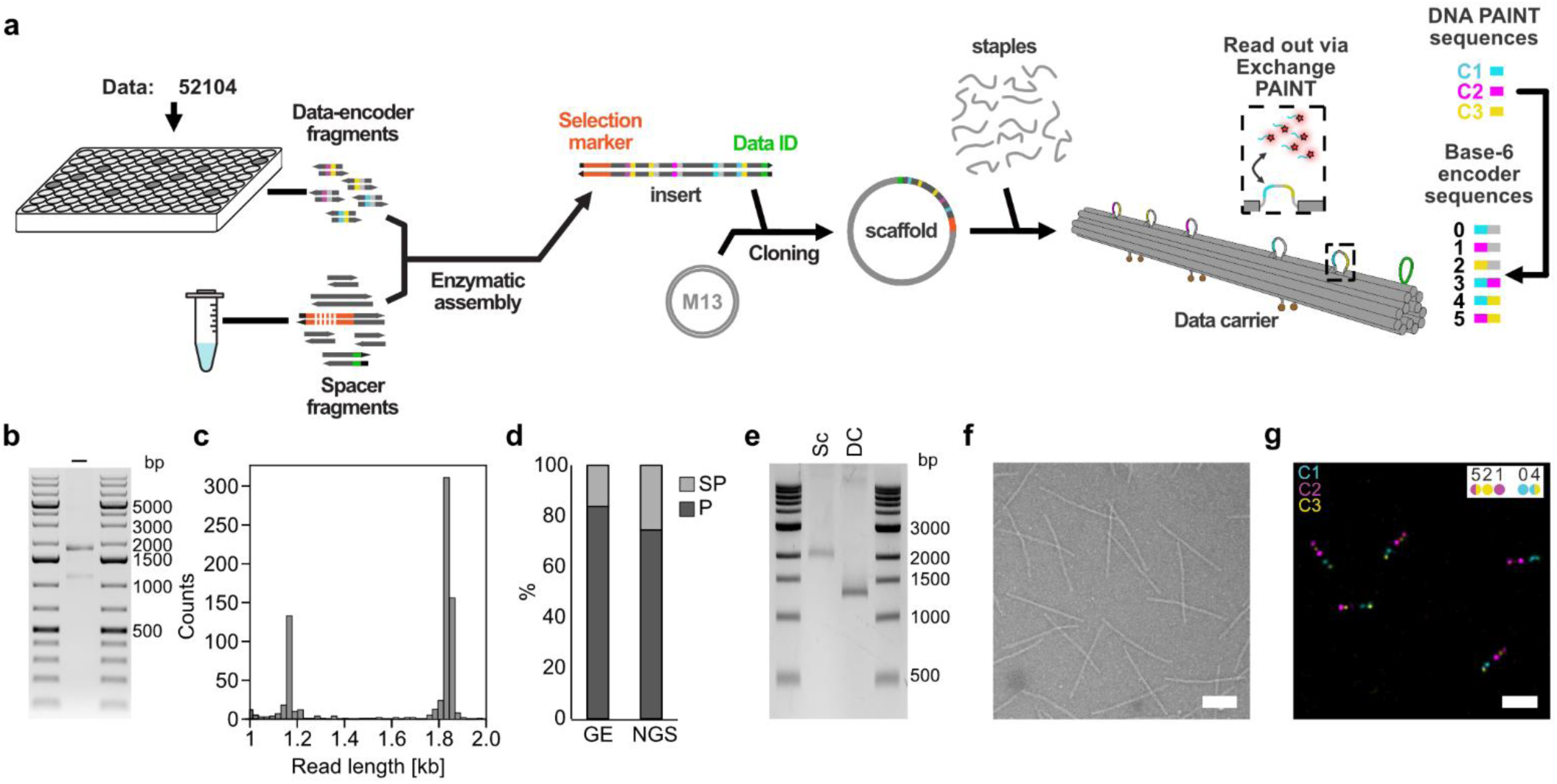
Combinatorial encoding of senary information into origami scaffold molecules with DOCS. **a,** Schematic of the workflow using DOCS to encode senary information. **b,** Agarose gel electrophoresis of the insert constructed with combinatorial, enzymatic assembly. **c**, Read length distribution of the insert extracted from nanopore sequencing data. **d,** Plot showing the fraction of product (P) and side-product (SP) **e,** Electrophoresis of scaffold (Sc) and data carrier (DC) generated from the insert in 2% agarose gel. **f,** Transmission electron microscopy micrograph of the data carrier (scale bar = 100nm) **g,** Exchange-PAINT image of the data carrier with the color codes for the different channels indicated and the color code of the encoded senary string (scale bar = 200nm).

To assemble the inserts, we created variants of DEFs with flanking regions that determine its position in the assembled insert. A collection of spacer fragments with flanks that overlap with the variant DEFs allow them to be assembled in a one-pot reaction. In addition, a selection marker was added to the assembly, to allow for selection after the insert was cloned into the M13 vector. (Fig1.a, Fig.S1).

To test the DOCS production pipeline, we first encoded a simple senary string with five different DES strands “52104”. The corresponding data carrier contained imaging sites from all three Exchange-PAINT channels used. We were able to construct the insert with high purity despite the complexity of the multicomponent assembly reaction, as confirmed with gel electrophoresis (~84% purity) and next-generation sequencing (~75% purity) (Fig.1b-d). The detected side-product produced by the assembly reaction, containing a truncated selection marker (Fig.S3), was later eliminated during the cloning step. This was confirmed by the homogeneously pure scaffold we produced from the insert (Fig.1e) that we successfully folded into the DOCS probe as characterized by TEM (Fig. 1f) and Exchange PAINT (Fig.1 g, Fig. S4). Processing of the Exchange PAINT data showed acceptable error rates (Fig. S5) for us to proceed to test the system for encoding meaningful data.

### Encoding texts with DOCS

To encode texts with DOCS we first converted the text into a senary string using Huffman encoding. The senary string was then divided into substrings of length 5, where each substring overlapped with 4 of the characters of the previous substring. These can be represented as a graph where nodes correspond to overlapping 4-character substrings of the complete senary string and the edges of the graph are represented by the individual DCs (Fig.2a, Fig. S6). Message extraction followed the reverse logic: first all DCs were converted into a graph (analogous to de Bruijn Graph assembly^26^ used in sequencing analysis), then the obtained string was decoded (Fig. S7). Following this strategy, we encoded a 23.3 bit (9 senary character) string (“stack”) using DOCS. The resulting scaffold library folded with high efficiency into the DC library (Fig.2b) that we further characterized using Exchange PAINT (Fig.3c, Fig. S8) where we detected all the constituent DCs of the library (Fig.2c). While we were able to successfully decode the message stored in the library, the probe frequency was not equal across all probes but followed a power law-like distribution (Fig. S9). Taken together with the error rates we observed, this would make the encoding of larger data unmanageable, so we investigated if we could implement error correction in the system. We tested two strategies: algorithmic error correction (AEC) and physical error correction (PEC).

**Fig. 2:**
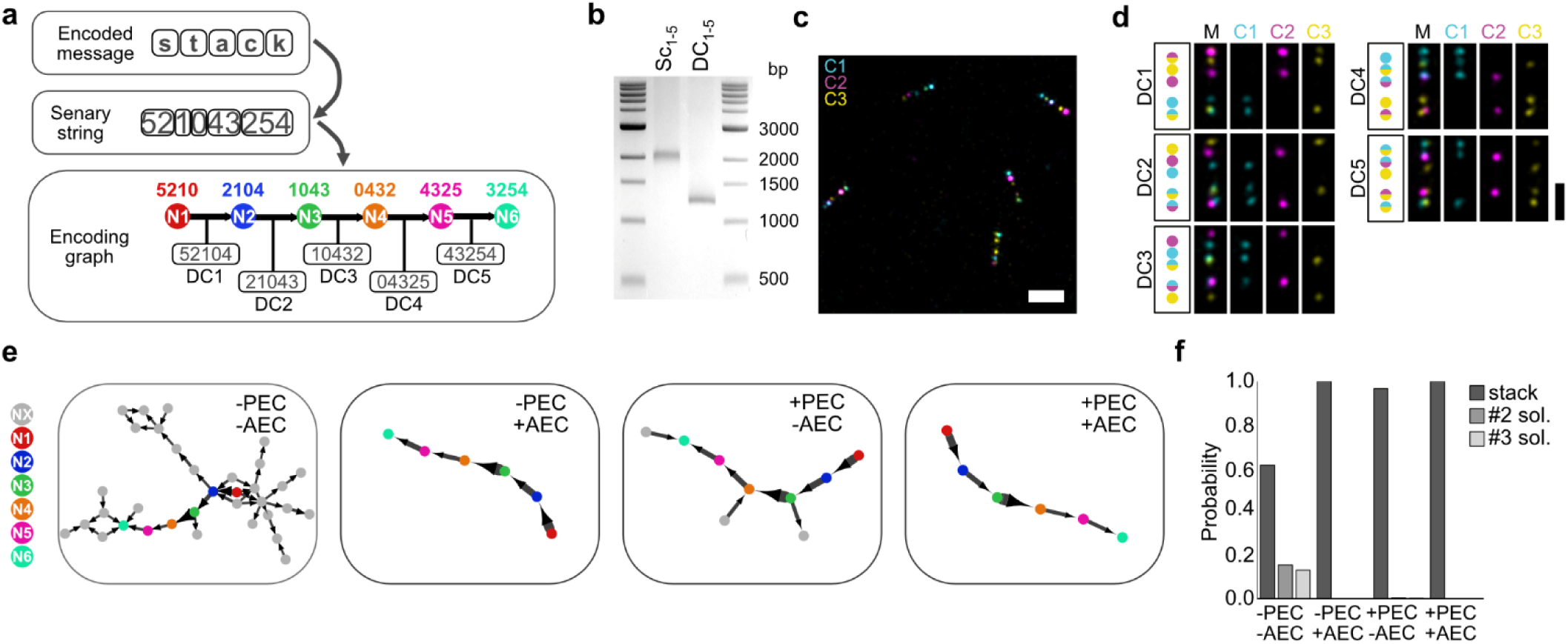
Encoding texts with DOCS. **a,** Schematic representation of the encoding strategy used to store text information in data carriers: text is converted to a senary string using Huffman encoding. The senary string is then converted into a graph by shifting a 4-character window through the senary string with a step size of one to generate the nodes of the graph. In this representation the data carriers (DC) constitute the edges of the graph. **b,** Agarose gel electrophoresis of the scaffold library (Sc1-5) and the DC library (DC1-5) encoding the text information. **c,** Exchange-PAINT image of the DC library (scale bar = 200nm). **d,** Colour codes and exemplary Exchange-PAINT images of the constituent DCs (DC1-5) of the library with images showing the merged (M) and separate channels (CH1-3) (scale bar = 100nm). **e,** The largest connected components of the graphs reconstructed from the decoded DC library using physical error correction (PEC) or algorithmic error correction (AEC) with the data encoding nodes (N1-6) colour highlighted along with non-intended nodes (NX, grey) originating from errors. **f,** Plot showing the calculated probability of the top three decoding solutions from the data recovered from the DP library using combinations of the different error correction strategies.

**Fig. 3:**
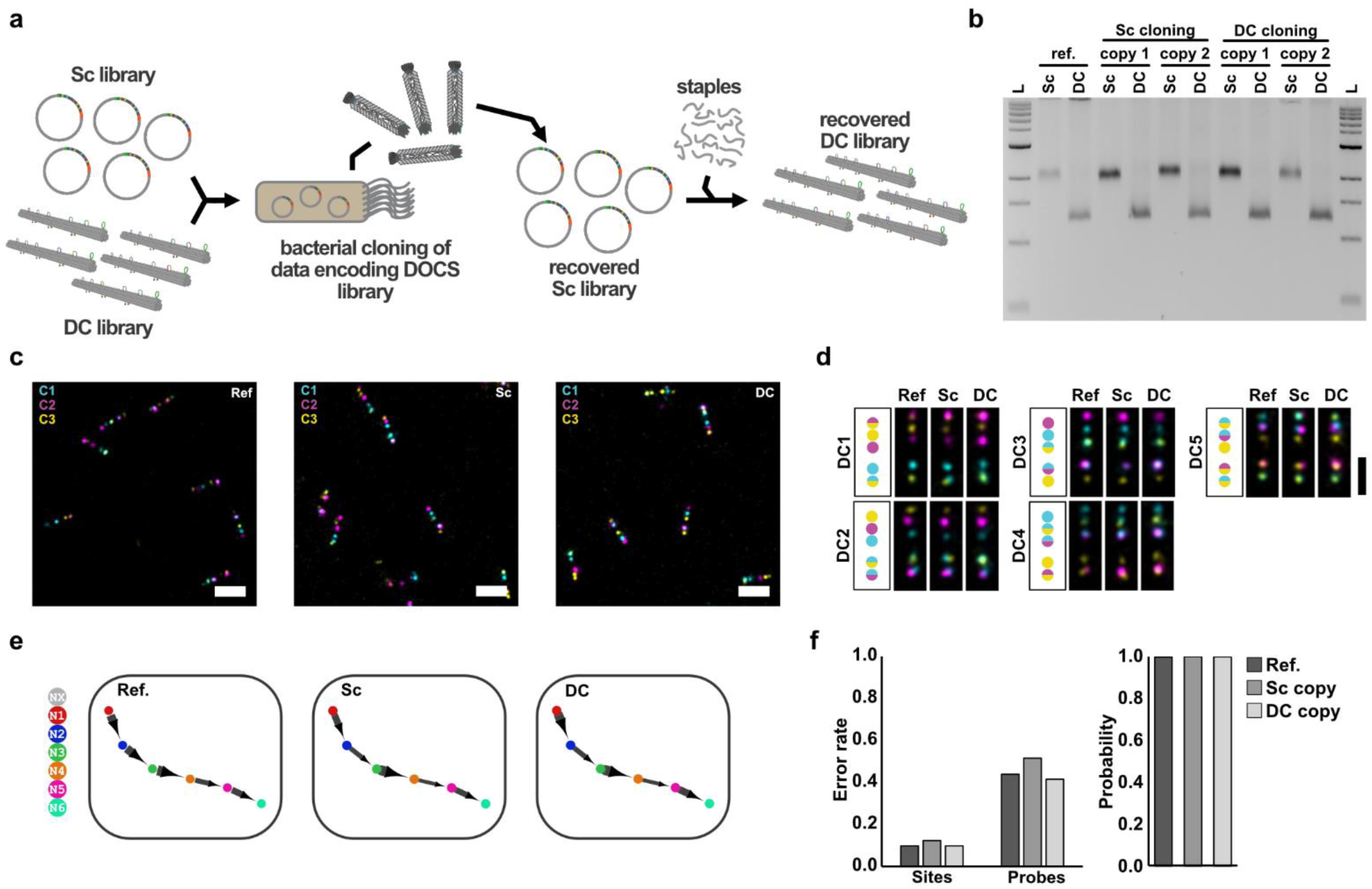
Molecular cloning of data encoded in DCs in bacteria. **a,** Schematic of the molecular cloning process of a data encoding DOCS library, where DH10B T1R E. Coli competent cells were transformed with either the scaffold (Sc) or data carrier (DC) library. **b,** Agarose gel electrophoresis of scaffold library (Sc) and DC library (DC) samples before and after cloning using the scaffold or the DC library as the source material for creating two independent copies in both cases. **c,** Exchange-PAINT images of DC library before and after cloning using the scaffold or the DC library as the source material (scale bars = 200nm). **d,** Colour code and exemplary Exchange PAINT images of the constituent data carriers (DC1-5) of the DC library detected in the source DC library and the copies originating from the scaffold library (Sc) or the DC library being used as source material (scale bar = 100nm). **e,** The largest connected components of the graphs reconstructed from the decoded data using AEC, obtained from the source DC library (left) and from DC library copies produced as the scaffold library (Sc, middle) or the DC library (DC, right) as source material. The data encoding nodes (N1-6) are colour highlighted along with gray nodes (NX) originating from errors. **f,** Plots showing the error rates observed per site and per probe (left) and the calculated probability of the intended information as the decoding solution using AEC (right) recovered from the source DC library and DC library copies produced using the scaffold library (Sc) or DC library (DC) as the source material.

AEC is an approach inspired by our observation that most incorrect DCs are closely related to their reference (Fig. S7). As part of the decoding pipeline, we kept only the highest frequency DCs from groups of closely related DCs for data reconstruction (Fig. S7). PEC was an experimental adjustment inspired by the fact that most errors were caused by imaging channel dropouts, which primarily affected mixed channel sites (Fig.S10), causing them to be read as single channel sites. Here, we utilized an extra imaging round to distinguish between real and erroneously identified single channel sites (Fig. S11). Implementing PEC resulted in an almost 10-times decrease in error rates (Fig.S11) and both approaches drastically reduced noise in the data reconstruction (Fig.2e) resulting in quasi-deterministic outputs (Fig.2f) showing the DOCS system can be scaled to larger datasets.

### Biological replication of DOCS-encoded data

One of the advantages of DOCS compared to classical DNA origami data storage approaches is that the information is sequence encoded. We therefore tested whether it is possible to biologically replicate the information stored in DOCS probes. For this, we transformed bacteria with the text-encoding DOCS library using the scaffold or the origami form of the library as input material (Fig.3a). We were able to produce new scaffold libraries from both inputs multiple times and fold them into DOCS probe libraries with similar efficiency as previously observed (Fig.3b). The copied libraries could be imaged with Exchange PAINT (Fig.3c, Fig.S12-13) and the constituent DCs of the libraries could be detected in all cases (Fig.3d). The frequencies of the DCs followed a similar distribution as observed before with some changes in relative frequencies of the constituent DCs in the library (Fig. S14). We were able to decode the message in all cases with almost deterministic certainty (P>0.99) using AEC (Fig.3e-f, Fig.S15). Furthermore, we observed largely similar error rates for the copied libraries with only the scaffold derived copy showing slightly higher error rates (Fig.3f).

### Thermostable data storage in DOCS

Conventional DNA origami structures typically unfold in a matter of minutes at temperatures above 60°C^27^ and only treatment with UV^28^, enzymatic crosslinking^29^ or heavy metals^30^ can stabilize them at higher temperatures. While these strategies preserve structural conformation, they may fail to protect the information-storing components or maintain compatibility with the detection techniques required for DNA origami data storage systems. Furthermore, we reasoned that information retrieval could still be done even by refolding the structures or perhaps from partially unfolded structures, as one only needs to be able to recognize the sequence of five signals. We therefore set out to test how resilient DOCS is to high temperatures.

For this, we exposed the text encoding DC library to temperatures ranging from 60°C to 80°C for 15 minutes to an hour in three different physical formats: suspended in origami storage buffer, as a dried pellet or absorbed on filter paper (Fig.4a). In liquid format the DC library stayed stable up until 60°C (Fig.4b, Fig.S16), while in dried pellet form, produced through PEG precipitation, we observed an aggregation that already occurred at 50°C (Fig.4b, Fig.S16). In comparison, when we stored the library absorbed on filter paper, it remained intact in all conditions (Fig.4b, Fig.S16). We also tested whether we could recover the disassembled structures by rerunning the temperature ramp we used for folding the DCs, which we were able to do for samples stored in liquid (Fig.4c) and dried form (Fig.S17). We then tested whether we could recover the information that was stored in the DCs and imaged two sets of samples: the DCs incubated at 80°C for 1h on filter paper (80°C FP), and the DCs incubated at 80°C for 1h in storage buffer and then refolded (80°C RF) (Fig.4d, Fig, S18-19). We were able to detect all the constituent DCs of the DC library for both samples (Fig.4e) with largely unchanged relative frequencies (Fig. S20) and were able to recover the stored information from them as well (Fig. 4f). The high temperature samples had slightly higher error rates, but similarly high data reconstruction probability as before (Fig.4g).

**Fig. 4:**
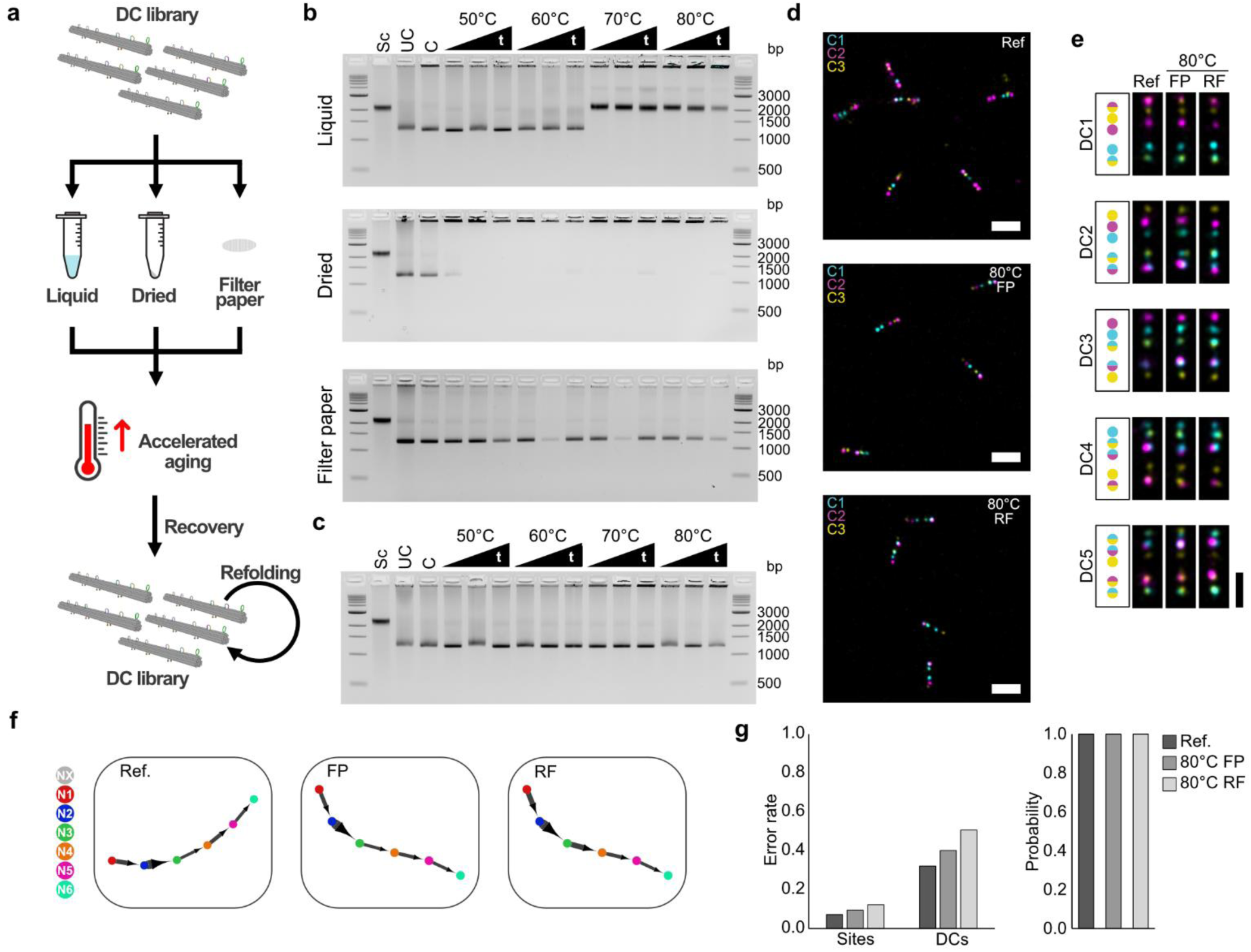
Thermostability of the DC library. **a,** Schematic of the workflow for the temperature stability experiment, where data-encoding DC libraries were incubated at elevated temperatures in different formats: liquid, dried pellets, or absorbed onto filter paper. The stability of the libraries in these formats was assessed after resuspending into liquid and refolding the sample if necessary. **b,** Agarose gel electrophoresis of DC library (DC1-5) samples incubated at 50-80**°**C for 15-60 minutes in liquid, solid pellets and absorbed onto filter paper along with scaffold library (Sc), uncleaned (UC) and cleaned control DC library (DC1-5) samples and DNA ladder (L). **c,** Agarose gel electrophoresis of DC library (DC1-5) samples incubated at 50-80**°**C for 15-60 minutes in liquid and refolded afterwards along with scaffold library (Sc), uncleaned (UC) and cleaned (C) control DC library (DC1-5) samples. **d,** Exchange-PAINT images of DC library before (Ref) and after aging at 80**°**C for 60min on filter paper (FP) or in liquid and recovered through refolding (RF) (scale bar = 200nm). **e,** Colour code and Exchange PAINT images of the constituent DCs (DC1-5) of the DC library detected in the reference DC library (Ref) and in the samples aged at 80**°**C for 60min on filter paper (FP) or in liquid and recovered through refolding (RF) (scale bar = 100nm). **f,** The graphs reconstructed from the data decoded using both AEC and PEC from the reference DC library and from DC library samples aged at 80**°**C for 60min on filter paper (FP) or in liquid and recovered through refolding (RF) with the data encoding nodes (N1-6) colour highlighted along with non-intended nodes (NX) originating from errors. **g,** Plots showing the error rates observed per sites and per DCs (left) and the calculated probability of the intended information as the decoding solution (right) recovered from the reference DC library and DC library samples aged at 80**°**C for 60min on filter paper (FP) or in liquid and recovered through refolding (RF).

### Random access and scalable data storage with DOCS

To test the implementation of random-access in the DOCS system we have encoded a 67-bit (26 senary character) message (“*docs stores info*”) across three sub-libraries (W1-3) (Fig. S21), each distinguished by a unique ID-sequence (Fig.5a). Following the characterization of the library with gel electrophoresis, electron microscopy and Exchange PAINT (Fig.5b-e, Fig. S22), we successfully recovered the error-free message using PEC (Fig.5f-g), which removed most non-intended DCs from the data and resulted in the intended DCs almost exclusively having the highest observed frequencies (Fig. S23). While AEC could further improve the probability scores of W2 and W3, it introduced an error in W1 by removing W1-DC3 due to its similarity to W2-DC5 (Fig.5f-g). We subsequently selected W3 sub-library using magnetic beads (Fig.5h). Exchange-PAINT imaging of the selected sub-library (Fig.5i, Fig.S24) confirmed the presence of constituent DCs (Fig.5j) and allowed for the recovery of the encoded information (Fig.5k) with low error rates and high selectivity (Fig.5l, Fig.S25).

**Fig. 5:**
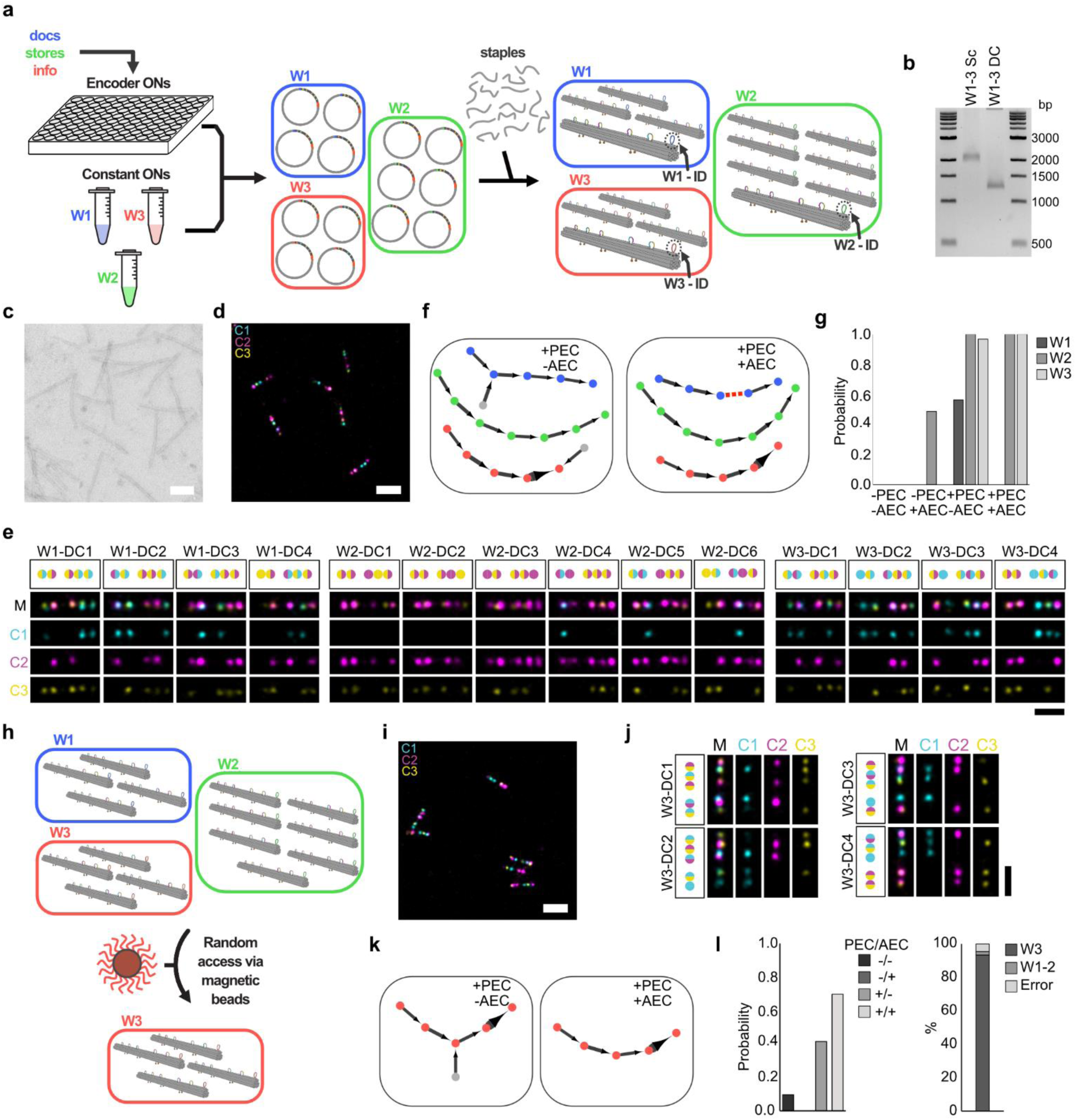
Multi word text encoding and random access with DOCS. **a,** The process used to encode multi word text information in data carriers: The individual words in the text are encoded into separate graphs and the data carriers (DC) representing the nodes in these graphs are constructed as described before, each set of DCs using a set of Constant ONs that carry an uniquely addressable ID sequence (W1-3 -ID) that is displayed as a loop on the folded DCs. **b,** Agarose gel electrophoresis of the scaffold library (W1-3 Sc) and the DC library (W1-3 DCs) encoding the multi word text information. **c**, Transmission electron microscopy micrograph of the DC library (scale bar = 100nm) **d,** Exchange-PAINT image of the DC library (scale bar = 200nm). **e,** The color codes and representative DNA PAINT images of the constituent DCs of the library with images showing the merged (M) and separate imaging channels (C1-3) (scale bar = 100nm). **f,** The graphs reconstructed from the decoded DC library using physical error correction (PEC) with and without algorithmic error correction (AEC) with the data encoding nodes of the different words colour highlighted (W1: blue, W2: green, W3: coral) along with non-intended nodes (grey) originating from errors. **g,** Plot showing the estimated probabilities of the intended information being the correct solution for the different words encoded in the library reconstructed using combinations of the different error correction strategies. **h,**The process of using magnetic beads carrying with capture DNA sequences complementary to the target word ID sequence for performing random access of the target word in the DC library. **i,** Exchange-PAINT image of the selected W3-DC library (scale bar = 200nm). **j,** The color codes and Exchange PAINT images of the constituent DCs of the selected W3-DC library (scale bar = 100nm). **k,** The graphs reconstructed from the decoded selected W3-DC library using physical error correction (PEC) with and without algorithmic error correction (AEC) with the data encoding nodes colour highlighted (coral) along with non-intended nodes (grey) originating from errors. **li,** Plots showing the estimated probabilities of the intended information being the correct solution for the data extracted from the selected W3-DC library reconstructed using combinations of the different error correction strategies (left) and showing the fraction of data extracted from the selected W3-DC library that encodes the selected word (W3), the other words (W1-2) in the full library or originating from error (right).

To explore the scalability of DOCS, we simulated storage of texts ranging from single words to full books up to 411 kilobytes (kB) in size (Table S1). As Huffman encoding’s uses the same DCs to encode recurring sections in texts, many DCs would be reused several times, resulting in encoding graphs with loops and competing solutions (Ext Fig. 1a). Although this could be partially alleviated by increasing the number of channels or sites (Ext Fig. 1b), it ultimately limited scaling to large texts. We therefore investigated the use of zlib compression^31^, reasoning that compression should remove redundancy and thus minimize the number of loops in the encoding graphs. Although the compression requires ASCII-encoded texts which adds overhead, especially for shorter texts, the benefits of compression outweigh the overhead for larger texts (Ext. Fig. 1c). Furthermore, the use of zlib-encoded binary data means that DOCS could be used to encode any data, not only texts.

The simulations were performed in triplicate using the power-law distribution of DC counts and error rates we extracted from the data (Fig. S26). Using a system with the same capacity as our experimental one and not using AEC (c=3, s=5) we could decode only short texts with low relative probability (Fig. S25). However, by increasing the number of sites and channels and using AEC we were able to decode all zlib-encoded texts with high relative probability (>=0.9) (Ext Fig. 1d). In addition, we found that we could reduce the number of required DCs by decreasing the overlap between adjacent DCs, effectively increasing the encoded information per structure. A shorter overlap required more sites or channels per DC for successful decoding, providing a tradeoff between information density per DC and total number of required DCs (table S2).

Compared to zlib-encoded texts, Huffman-encoded texts typically resulted in graphs with more branching points (Ext Fig. 1e-f), which prevented us from decoding larger Huffman encoded texts even when no errors were added (Ext Fig. 1g). In the presence of errors, the use of AEC proved necessary. However, applying AEC frequently resulted in the removal of correct, reused DCs in the data, leading to the breaking of the encoding graph into multiple components and thus ultimately preventing decoding (Ext Fig. 1h). Again, using zlib-encoding and a higher number of channels and sites fixed this and resulted in successful decoding (Ext. Fig. 1g-h).

### Creating Molecular authentication system with stochastic DOCS

Beyond deterministic information storage, the large combinatorial space of the DOCS platform can be leveraged to create a molecular authentication system that satisfies the fundamental requirements of a Physical Unclonable Function (PUF)^32^. In simple terms, a PUF deterministically yields an output signal (response) for a specific input signal (challenge). However, since the generation of the system’s components and their association relies on stochastic physical processes, the precise mapping between any given challenge and response remains completely unpredictable. To implement this, using building blocks derived from the DOCS system (Fig.S27-28) we generated a library of random inserts (corresponding to the response of the PUF) in a one-pot reaction and linked random ID molecules—acting as stochastic seeds (corresponding to the challenge of the PUF)—to each unique DOCS insert (Fig. S29). From this mixture, we generated a scaffold library and subsequently folded a DC library, where each individual DC serves as a distinct physical PUF instance (Fig. 6a).

**Fig. 6:**
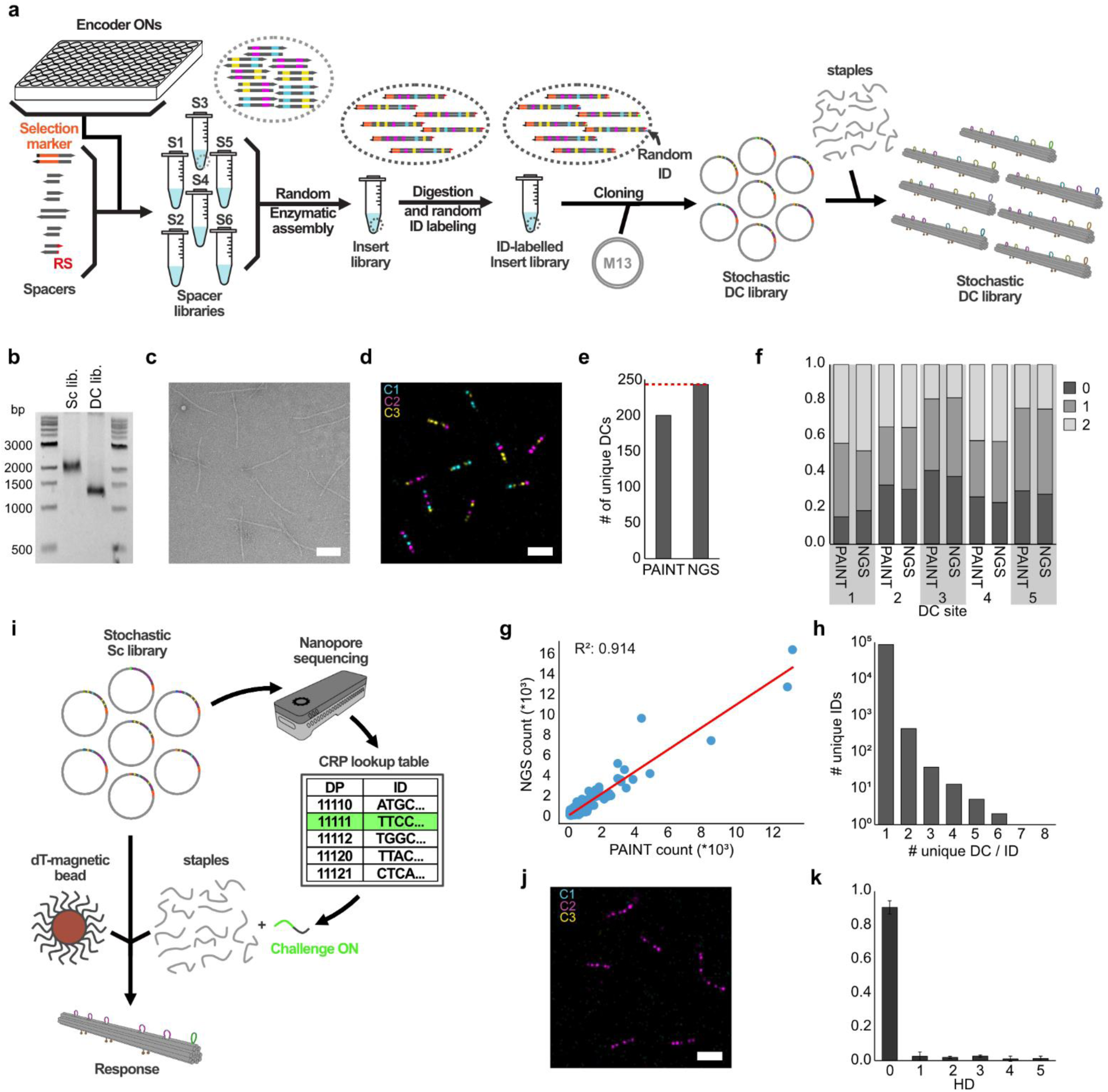
Creating a molecular authentication system with stochastic DOCS. **a,** The workflow for generating stochastic, three character DOCS library with randomly associated ID sequences: Spacer libraries are generated separately using spacer ONs and associated encoder ONs, then the spacer libraries are used to create the insert library in a one-pot reaction, inserts are labelled using a random ID library subsequently and are cloned into M13 to create the stochastic 3 character scaffold library, which then can be folded into the DC library using a single set of staples. **b,** Agarose gel electrophoresis of the 3-character scaffold library (Sc lib.) and the DC library (DC lib.). **c,** Transmission electron microscopy micrograph of the DC library (scale bar = 100nm) **d,** Exchange-PAINT image of the DC library (scale bar = 200nm). **e,** Plot showing the number of detected unique DCs by Exchange PAINT measurement of the DC library (PAINT) and Nanopore sequencing of the DC library (NGS) with the red dashed line indicating the size of the complete library. **f,** Plot showing the relative frequencies of the different symbols per site on the DCs measured using DNA PAINT imaging of the DC library and Nanopore sequencing of the scaffold library. **g,** Plot showing the sequencing read counts versus the confidence-weighted probe counts in the PAINT data set for the unique DCs detected in the library with the linear regression fitted to the data (red line). **h,** Plot showing the number of unique DCs associated with unique ID sequences. **i,** The workflow for mapping the random ID sequences to specific DCs in the library to create a challenge-response-pair (CRP) lookup table and performing and authentication test: The scaffold library was sequenced and the lookup table is generated with associated unique DCs and ID sequences then a Challenge ON complementary to the target ID sequence and to the poly-T ONs displayed by the magnetic beads is used for the authentication test via solid state synthesis approach^46^. **j,** Exchange-PAINT image of the response of the authentication test sample (scale bar = 200nm). **k,** Plot showing the relative frequency of probes detected in the response DC samples with codes the stated Hamming distance from the intended DC’s code.

The resulting stochastic DC library folded with high efficiency, as demonstrated by gel electrophoresis (Fig. 6b), electron microscopy (Fig. 6c), and Exchange-PAINT imaging (Fig. 6d, Fig.S30). To complement the optical readout, we further characterized the library using Nanopore sequencing. In a single Exchange-PAINT field of view (FOV), we detected over 80% of the theoretically possible codes (Fig. S31), while the NGS dataset confirmed the presence of all possible codes (Fig. 6e). The site-wise symbol distribution was largely uniform and showed a high degree of correlation between the PAINT and NGS datasets, confirming the unbiased nature of the library synthesis (Fig.6f).

To quantify the impact of synthesis bias on the system’s security performance, we calculated the Empirical Hamming Distance (weighted mean HD) of the library. The results showed negligible bias, with the empirical distance reaching 96.4% of the theoretical maximum (Fig. S32). Furthermore, the abundance of unique DCs was strongly correlated between the PAINT and NGS datasets. This reinforced the genotype-phenotype link inherent to the platform (Fig. 6g) and allowed us to generate a robust Challenge-Response Pair (CRP) lookup table linking each unique DC optical code to its specific genetic ID sequence. Analysis confirmed that the vast majority of unique ID sequences were associated with a single unique optical code (1.01 ± 0.10), with approximately 360 unique ID sequences mapping to each unique DC on average (Fig. S32).

Encouraged by these metrics, we validated the system experimentally by performing a magnetic bead selection on the library using a challenge oligonucleotide derived from the CRP lookup table (Fig. 6i). The physical response of the library was highly specific (Fig. 6j, Fig. S33): over 90% (0.90 ± 0.04) of the detected DCs displayed the anticipated code (Fig. 6k). This resulted in a low bit error rate of 0.05 ± 0.01, corresponding to a system reliability of 95%.

## Discussion

While DOCS overcomes many of the limitations inherent to previous DNA origami-based approaches it still faces bottlenecks when considered for implementation on a practical storage scale.

One of these is the synthesis strategy. Although the steps in this proof-of-concept implementation were carried out manually, the larger part of the workflow (PCR assembly, PCR cloning, and bead-based purification) is highly automatable using commercially available liquid-handling robots. Integrated biofoundry solutions capable of automating the entire workflow, including the bacterial production steps, already exist and could be adapted to rapidly scale DOCS synthesis^33^.

Secondly, while Exchange-PAINT^34^ as a read modality offers several advantages, such as parallel readout and low consumable costs, the current data acquisition protocol is not scalable. The primary limitation is the prolonged data acquisition time, which could be reduced by several orders of magnitudes utilizing recently developed strategies for accelerated DNA PAINT imaging^35–37^. Furthermore, the sophistication and cost of the required instrumentation (a complete TIRF microscope setup) can also be drastically minimized^38^. Additionally, the multiplexing capacity of DNA-PAINT can be expanded from the three channels used here to ten^34^ or even thirty^39^ to increase the platforms encoding capacity. The path-finding algorithm can also be improved by taking inspiration from state-of-the-art genome assembly algorithms that use variable k-mer lengths and more efficient path pruning^40,41^. More efficient error handling would allow the data to be encoded with a lower number of channels and sites per probe, increasing the scalability. To summarize there are multiple established strategies to considerably improve the data collection strategy used in the system while retaining Exchange-PAINT, though transitioning to a non-super-resolution imaging approach may ultimately prove to be the most scalable path forward^42^.

Beyond data storage, physically unclonable functions (PUFs) have previously been implemented using DNA in vivo^43^, in vitro (chemical unclonable functions (CUF) by Luescher et al.^44^), and via DNA origami structures (PartiPUFs by Dass et al.^45^). While the primary application of DOCS presented in our work is data storage, our DOCS-based, proof-of-concept molecular authentication platform demonstrates distinct advantages over these existing systems.

When evaluating system capacity, the size of the challenge-response pair (CRP) space in DOCS is significantly larger than what PartiPUF allows, though smaller than the vast sequence-based response space of CUF. However, this is mitigated by the theoretical size of the DOCS challenge space (~10^15^). Static optical labels like PartiPUF possess enormous theoretical instance capacities but offer limited CRP space per instance. Conversely, both DOCS and CUF function as Strong PUFs, capable of processing massive challenge spaces from a single physical pool.

In terms of security, while PartiPUF labels are inherently physically unreplicable, chemical pools like CUF must implement complex mitigations to prevent copying via enzymatic amplification. In the case of DOCS, while the underlying scaffold library can be copied via molecular cloning, reproducing the DNA alone does not replicate the functional system. The precise combination of DNA staples acts as a necessary physical encryption key; without them, no meaningful response can be achieved. Furthermore, while the current DOCS authentication implementation uses bead-based probe selection for its challenge mechanism, shifting to an in-situ fluorophore-labelled challenge strand would allow for non-destructive, repeated interrogations. Ultimately, the current limitations of the authentication system can be readily surmounted by simply increasing the complexity of the DOCS library, as described for the data storage application.

## Conclusion

With DOCS we have built a system that breaks away from the reliance of DNA origami-based data storage approaches on DNA hybridization to store information, while maintaining the same extent of modularity. This is achieved by using a combinatorial enzymatic strategy to encode information directly into the scaffold molecule. We have demonstrated that this approach allows the storage of text information in a quasi-deterministic fashion and facilitates the implementation of random access. The DOCS system has also proved to have an unprecedented thermostability and the capability of being copied via molecular cloning. Furthermore, we have shown through simulations that the primary obstacle to the system’s scalability, the effect of repeated fragments on the decoding process, can be mitigated with easily implementable measures. This includes reducing the redundancy via data compression or increasing the encoding capacity of DOCS by utilizing more imaging channels or data encoding sites per probe. Finally, we highlighted the combinatorial power of the platform by creating a DOCS-based molecular authentication system, which demonstrated uniform diversity and highly reliable challenge-response capabilities. Through these demonstrated capabilities, we have shown that DOCS provides an automatable, scalable, and robust approach to DNA origami-based data storage, utilizing a finite set of universal components to limit encoding costs and establish structural DNA nanotechnology as a highly competitive alternative to purely sequence-based approaches.

## Supporting information

Supplementary_file_sequences

## Figures

**Extended Fig. 1.**
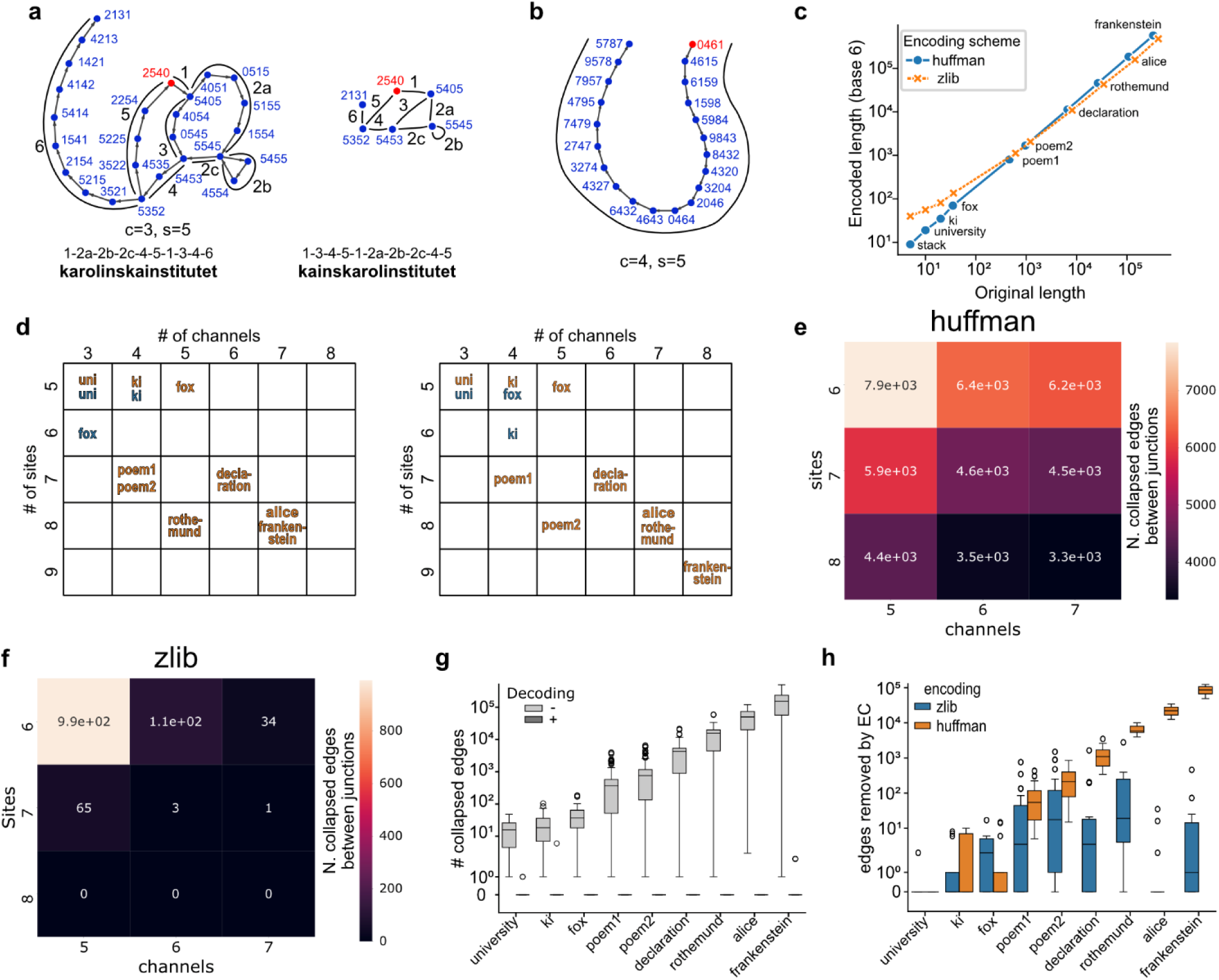
Increasing the number of channels and sites per probes allows for storage of kilobytes of data in simulations. Successful decoding means the correct path is found with a relative probability >= 0.9. **a,** Left: graph of Huffman-encoded text “karolinskainstitutet”. Right: Same graph with compressed edges. Red node is the starting node of the correct path. **b,** As (**a**), but encoded with an additional channel. **c,** Lengths of original texts and texts encoded with senary code. **d,** Overview of lowest channel number/site number combination that allowed successful decoding in simulations. Left: no added errors; Right: with physically corrected errors, and after applying algorithmic error correction. **e-f** Number of collapsed edges between branch points in graphs obtained from encoding “rothemund” text with Huffman (**e**) or zlib (**f**), without errors, with various numbers of channels and sites. **g,** Distribution of numbers of collapsed edges between branch points in graphs without errors or error corrections that were successfully decoded (+) or not (−). **h,** Distribution of numbers of edges removed by algorithmic error correction in simulations without errors.

**Table S1.**
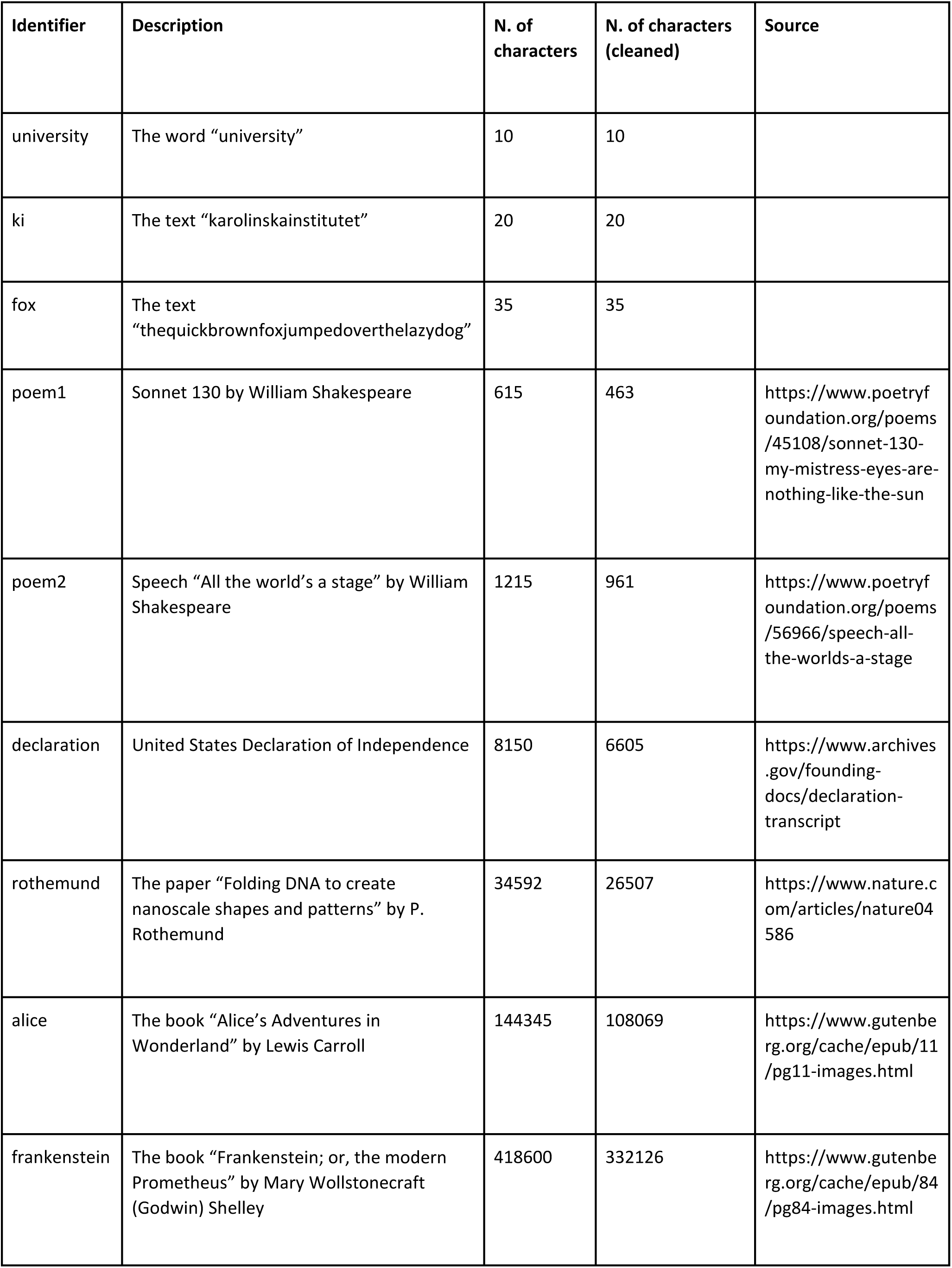
Texts used in simulations. Cleaned texts were used for Huffman encoding and only contained lowercase letters without punctuation or spaces.

**Table S2.** Increasing the shift reveals a tradeoff between total structures and structure signals. The text “alice” (Table S1) was encoded using shifts of 1 to 4. Increasing the shift lowers the amount of required structures but requires either more signals per structure or more color channels to compensate. Success was defined as having the correct path be identified with relative probability >= 0.9 compared to all other paths.

| Shift | Different unique structures (no errors) | Lowest successful color/kmer combination (no errors) | Lowest successful color/kmer combination (corrected errors) |
| --- | --- | --- | --- |
| 1 | 78 965 | C7, K8 | C7, K8 |
| 2 | 39 484 | C7, K9 or C8, K8 | C7, K9 or C8, K8 |
| 3 | 26 323 | C7, K9 | C8, K9 |
| 4 | 19 743 | C8, K10 | C9, K10 |

## Methods

### Design of oligos used for DOCS

The insert was designed to contain a selection marker (Kanamycin resistance gene cassette), five data encoding sequences (DES) flanked by spacer sequences (spacer fragments (SF)) and a 25nt long ID sequence used for random access. The SFs were designed to be 84nt long, except for SF4, which is 168nt long, placing the DES in distances easily resolvable by Exchange PAINT and introducing an inbuilt asymmetry aiding the code determination of data carriers. Each DES contains two pairs of Exchange PAINT docking sites flanked by 3nt polyT spacers. For DES containing a single type of Exchange PAINT docking site, a pair of “blank” docking sites were used to keep the DES length constant. Data encoder fragments (DEF) used for the assembly of the inserts were designed to contain the DES flanked by 19nt sequences complementary to adjacent SFs.

### Production of inserts

All oligonucleotides were ordered from Integrated DNA Technologies. The Kanamycin resistance cassette and spacer fragment one fusion product (KANr-SF1) was ordered as a gBlocks gene fragment the rest of the SFs and their complements (SF2-SF6, SF2c-SF6c) were ordered as PAGE Ultramers, while PCR primers and the EONs and their complements were ordered as standard desalted oligos (see Supplementary_file_sequences). Double stranded KANr-SF1 was produced by 30-cycles of PCR amplification using flanking primers (KANr-SF1 FW: GGAAACAGCTATGACCATGATTAC, KANr-SF1 REV: GTTGCGCAAACTATTAACTGGCG) and Phusion High-Fidelity DNA Polymerase (New England Biolabs, M0530L) followed by a subsequent clean up using Monarch PCR & DNA Cleanup Kit (New England Biolabs, T1030L) following the manufacturer’s instructions. The concentration of the double stranded KANr-S1 stock was determined using a Nanodrop 2000 instrument (Thermo Scientific). DOCS insert molecules were assembled in parallel PCR reactions containing 15 nM of each double stranded SF, 75 nM of each double stranded EON in the set of EONs belonging to the particular insert, Phusion High-Fidelity DNA Polymerase (New England Biolabs, M0530L), 1X GC Buffer, 200μM dNTP in 10 μL reaction volumes using 30 PCR cycles. The assembled inserts were amplified in a PCR reaction containing 5 μL of the assembly reaction, 10 μM of flanking primers (DOCS insert FW: GGAAACAGCTATGACCATGATTAC, DOSC insert REV: GCAGGTCGACTCTAGAGGATCC), 1X GC Buffer, 200μM dNTP and Phusion High-Fidelity DNA Polymerase (New England Biolabs, M0530L) in 50 μL reaction volumes using 30 PCR cycles. DOCS insert molecules were then purified in parallel using 1.8x sample volume of AMPure XP beads (Beckman Coulter, A63881) following the manufacturer’s instructions and the 4200 TapeStation System (Agilent) was used with D5000 ScreenTapes (Agilent, 5067-5588) to inspect them and determine their concentration.

### Production of scaffold libraries

Linearized M13mp18 RFI (New England Biolabs, N4018S) was produced through restriction digestion using BamHI HF (New England Biolabs, R3136S) and EcoRI HF (New England Biolabs, R3101S) followed by purification using Monarch PCR & DNA Cleanup Kit (New England Biolabs, T1030L) following the manufacturer’s instructions. The concentration of the linearized M13 stock was determined using a Nanodrop 2000 instrument (Thermo Scientific). The DOCS inserts were cloned into the linearized M13 in parallel circular polymerase extension cloning (CPEC)^46^ reactions containing 140 pM linearized M13, 280pM DOCS insert, 1x GC Buffer, 200 μM dNTP and Phusion High-Fidelity DNA Polymerase (New England Biolabs, M0530L) in 20 μL reaction volumes using 30 PCR cycles. CPEC reactions were then pooled and 2 μL of the pool was transformed into MegaX DH10B T1R cells (Thermo Scientific, C640003) by electroporation at 1.5 kV using the Eporator instrument (Eppendorf). The transformation mixture was then immediately resuspended in 1 mL of Recovery Medium (Thermo Scientific) and incubated in a shaking incubator for 40min at 37 °C. The culture was then transferred into 500 mL of growth medium (500 mL of 2x Yeast extract Trypton (Sigma-Aldrich, Y2377), 1 mL of 50 mg/mL Kanamycin, 4.8 mL of 1M MgCl_2_, 100 μL of AntiFoam 204 (Sigma-Aldrich, A8311)) in Thomson’s Ultra Yield™ Flasks and was grown for 48h at 37°C. The scaffold library was extracted from the produced phages following the purification steps of the phage-amplification-in-bacteria protocol described in previous work (reference). The resulting scaffold library was further purified using PEG cleaning of scaffold PEG precipitation. The scaffold solutions were adjusted to 100 nM using 10 mM TRIS (pH 8.5) and were mixed 1:1 (v/v) with a PEG solution containing 20% PEG4000 (w/v) and 1 M NaCl. The solutions were briefly vortexed and centrifuged at 16,000 g at room temperature for 25 min using a microcentrifuge. The supernatant was carefully collected and a buffer exchange to 10 mM TRIS (pH 8.5) was performed by five rounds of ultrafiltration using 100kDa Amicon Ultra micro centrifugation filters (Millipore).

### Design and production of data carriers

The caDNAno software (cadnano.org) was used for designing the data carrier nanostructure consisting of twelve long parallel helices arranged in a honeycomb-lattice. The top central helix was designed to contain the unpaired, single-stranded loop regions displaying the sequences of the EONs and ID sequence. On the opposing face of the structure, three pairs of staple oligonucleotides were designed to have breakpoints facing “downwards” for surface immobilization sites modified with biotin. At the termini of the helices, staple oligonucleotides with 5-base long poly-A (A-caps) or 3-base-long poly-C (C-caps) extensions were used to prevent probe dimerization. All staple oligonucleotides were ordered from Integrated DNA Technologies standard desalted oligos (see Supplementary_file_sequences).

### Folding and purification of data carriers

Data carrier (DC) libraries were produced by folding reactions containing 10 nM Sc library, DC staple set (50 nM of each core staples and A-caps, 250 nM of each biotin staples) and on-tenth of the final reaction volume of folding buffer (100 mM MgCl_2_, 50 mM TRIS, 10 mM EDTA). The folding was carried out by an overnight temperature ramp that consisted of the following steps: heat denaturation at 65 °C for 15min, quick transition to 60 °C followed by slow cooling from 60 °C to 40 °C over 20 hours, then another quick transition to 20 °C. Excess of staple oligonucleotides were removed by repeated washing with storage buffer (identical to 1x folding buffer) in 100kDa Amicon Ultra micro centrifugation filters (Millipore) by concentration steps carried out by centrifugation at 14,000 g for 2 min. DC concentration was measured with the Nanodrop 2000 (Thermo Scientific). Quality assessment of the DCs was performed by running them in 2% agarose gel (0.5X TBE, 10 mM Mg2+, 0.5 mg/ml ethidium bromide) at 90 V for 2 h in ice bath.

### Transmission electron microscopy (TEM) imaging of DCs

Formvar/carbon-coated 400-mesh copper grids (FCF400-Cu, Electron Microscopy Sciences) were glow-discharged prior to use. A staining solution was prepared by adjusting the pH of 400 µL of 2% (w/v) uranyl formate with 8 µL of 1 M NaOH, followed by centrifugation at 16,000 × g for 5 minutes to remove precipitates. A 3 µL drop of purified DNA origami sample was applied to the grids for 90 seconds. Excess sample was removed by blotting with filter paper, and the grids were immediately stained with 15 µL of the uranyl formate solution for 40 seconds and blotted dry. Micrographs were acquired using a Talos 120C G2 transmission electron microscope (Thermo Fisher Scientific) operating at an accelerating voltage of 120 kV and a magnification of 57,000 × corresponding to a calibrated pixel size of 2.475 Å/pixel.

### Sample preparation for Exchange PAINT

Glass-bottom µ-Slide VI chambers (ibidi) were used for Exchange PAINT experiments. Channels were washed with 200μL of Buffer A (10 mM Tris-HCl, 100 mM NaCL, 0.05% Tween-20, pH 7.5) three times and were incubated with 100 μL of biotinylated-BSA (1 mg/mL in Buffer A, Sigma Aldrich) in Buffer A for 5 min. The channels were then washed with 200μL of Buffer A two times. Subsequently the channels were incubated with streptavidine (0.5 mg/mL, Thermo Scientific) in Buffer A for 5min. The channels were washed again with 200 μL of Buffer A two times. 100 μL of a concentrated solution of 80nm gold nanoparticles (Sigma Aldrich, 742023) resuspended in Buffer A was flushed into the channels and incubated for 5min followed by two washing steps with 200μL of Buffer A. The channels were then washed with 200 μL of Buffer B (5 mM Tris-HCl, 10 mM MgCl2, 1 mM EDTA, 0.05% Tween-20, pH 8) two times before structures resuspended in Buffer B (100 pM) were flushed in. The structures were then incubated for 10 min followed by washing with 200μL of Buffer B two times. The channels were then flushed with imaging buffer (40 pM Atto 550-labeled imager (C1=TATGTAGATC, C2= GTAATGAAGA, C3= CATACATTGA or C error= CTTATTGTAG (Integrated DNA Technologies)) (Table SX.) containing oxygen scavengers PCA (Sigma Aldrich, 37580) and PCD (Sigma Aldrich, P8279) and Trolox (Sigma Aldrich, 238813) at the previously described concentrations^34^. Channels were washed with 200 μL of Buffer B four times in between imaging rounds.

### Exchange PAINT imaging of data carriers and data pre-processing

Exchange-PAINT^34^ experiments were conducted with a microscope with Nikon Eclipse Ti-E microscope frame with the Perfect Focus system (Nikon Instruments) and an objective-type TIRF configuration using an iLAS2 circular TIRF module. A 1.49 NA CFI Plan Apo TIRF 100× Oil immersion objective was used alone or with an extra 1.5× magnification resulting in a final pixel size of 87nm. For illumination an OBIS 561 nm LS 150 mW laser was used with custom iLas input beam expansion optics (Cairn) optimized for reduced field SR imaging. A filter cube (89901, Chroma Technology) containing an excitation quadband filter (ZET405/488/561/640x, Chroma Technology), a quadband dichroic (ZET405/488/561/640bs, Chroma Technology) for cleaning up the illumination light and a quadband emission filter (ZET405/488/561/640m, Chroma Technology) was used in addition to an emission filter (ET595/50m, Chroma Technology) for filtering the collected light. An iXon Ultra 888 EMCCD camera (Andor) was used for recording. The Micromanager software was used for data acquisition using frame-transfer mode of the camera, 300 ms exposure time, 10 MHz readout rate and no EM gain and 9000 frames for each channel in a 512×512px size field of view. The Picasso software platform^47^ was used for the reconstruction of the microscopy data from the recorded movies using the MLE algorithm and the adaptive intersection maximization-based method (AIM) based drift correction^48^. Localizations were linked to form events using the software’s “Link localizations” feature and a final round of drift correction was used on individual sites selected by automated picking using the software’s “Undrift from picked” feature. Finally, channels were then aligned using the Picasso Render software’s “Align channels” feature.

### Exchange PAINT data processing

For each data set data collected from a single field of view was used. Events lying the non-overlapping part of the data sets of the different channels were removed. A low resolution filtered binary image was produced from the joint datasets using Otsu’s binarization in which the data carrier (DC) positions were detected by size-thresholded contour detection after a universal pixel dilation step. Events in the different channel datasets falling in the 260×260nm windows centered around the detected DC contours were extracted and grouped to belong to putative individual DCs (Fig. S34).

Events belonging to individual DCs were then screened and filtered: first DC site positions were identified with local maxima detection in high resolution renders of the events, and after keeping only the centrally clustered sites the positional identity of sites were determined from their pairwise distances for DCs with five detected sites and only DCs with sites showing high linearity and small deviation from the designed pairwise distances were kept. Finally, events in each channel are all assigned to DC site closest to them and filtered using DBSCAN clustering. The final DC site positions are recalculated using the mean position of events from all channels that are assigned to them (Fig.S35).

The identity for individual DCs was determined with a probabilistic approach using custom scripts as follows. Event number distribution per DC sites are calculated for each channel from the full dataset, which contain site assigned events from sites that are “ON” (generated by EONs with docking sites for the channel) and that are “OFF” (generated by EONs with no docking sites for the channel) in that given channel, as the DC site positions are determined from the merged data. These bimodal Poisson distributions are then fitted to extract λ_ON_ and λ_OFF_ (the event frequencies of the “ON” and “OFF” sites respectively) and *P*(*C_i_* = *ON*) and *P*(*C_i_* = *OFF*) (probabilities of observing an “ON” and “OFF” sites in the channel in the data) for each channel (Fig.S36). For each site in each individual DC the number of events in the given channel (*n_i,j_*) is used to calculate *P*(*n_i,j_*|*C_i,j_* = *ON*), the probability of observing the detected number of events in channel *i* in position *j* given that channel *i* in position *j* is being “ON”, using the following Poisson formula:

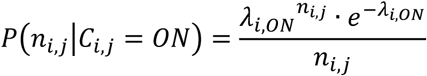

and *P*(*n_i_*|*C_i_* = *OFF*) correspondingly:

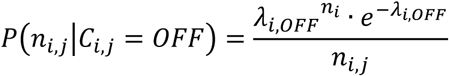

Using these probabilities *P*(*C_i,j_* = *ON*|*n_i,j_*), the probability of that channel *i* is being “ON” in position *j* given the detected number of events in channel *i* in position *j*, is calculated for each channel in each position using the following Bayesian probability formula (Fig.S36):

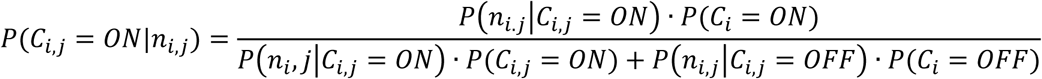

Finally, *P*(*C_i,j_* = *OFF*|*n_i,j_*) is calculated as 1 − *P*(*C_i,j_* = *ON*|*n_i,j_*).

Using these conditional probabilities for each channel the probabilities of each symbol being the identity of a given DC position is calculated as the product of the corresponding probabilities, for example (Fig.S36):

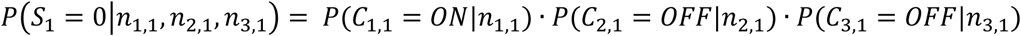

All non-intended states (e.g. all channels being “ON”) are assigned to a common symbol indicating an ambiguous position (“X”). In the case of data sets using physical error correction (PEC), all states with a single channel being “ON” but the error correction channel being “OFF” are also assigned ambiguous.

Finally, the probability of a given set of symbols being the code of an observed DC is calculated from the probabilities of positional symbol identities, for example:

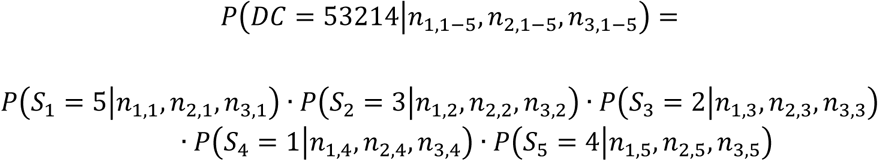

All probabilities of all possible sets of symbols are calculated for each DC, and the highest one is assigned as the DC identity (Fig.S36). For data reconstruction only DCs not containing ambiguous positions and with P>0.99 or P>0.9 for the datasets using PEC are kept with probability weighted counts.

### Recloning of scaffold and DC library

2μL of 1nM scaffold or DC library was transformed into MegaX DH10B T1R cells (Thermo Scientific, C640003) by electroporation at 1.5 kV using the Eporator instrument (Eppendorf). The recloned scaffold library was produced from the transformed cells as described earlier.

### Measuring thermostability of DC library

DCs were folded as described earlier to result in a final 10nM origami concentration. The thermostability measurements were performed on 100 µL aliquots from pooled 10 nM origami folding reactions. The structures were first purified and pelleted by one round of PEG precipitation^49^.

For the heat treatment on DCs in solution, the pellet was left to dry for 5 minutes and resuspended in 100 μL of storage buffer for 1 hour. The samples were incubated at 50 °C, 60 °C, 70 °C or 80 °C for 15 min, 30 min and 1 hour, and immediately analyzed by gel electrophoresis. The remaining volumes were then re-folded following the original folding program and analyzed by agarose gel electrophoresis and TEM.

For the heat treatment on filter paper, the pellet was left to dry for 5 minutes and resuspended in 50 μL storage buffer for 1 hour. For each condition, two samples were successively pipetted onto a 1 cm diameter circle of Whatman filter paper and briefly dried at 37 °C between additions. The filter papers were then placed in open Eppendorf tubes and incubated in an oven for the respective incubation durations and temperatures. After heating, the papers were rehydrated in 100 μL storage buffer overnight at room temperature on a shaker at 500 rpm, and the samples were analyzed by agarose gel electrophoresis and TEM.

For the heat treatment on dried structures, the pellet was left to dry for 15 minutes, before incubating for the respective durations and temperatures. The samples were then resuspended in 100 μL storage buffer overnight at room temperature on a shaker at 500 rpm and analyzed by gel electrophoresis. The remaining volumes were re-folded following a folding program starting at 80 °C and analyzed by agarose gel electrophoresis.

### Random access of information stored in DC library

Scaffold library encoding the three randomly accessible scaffold sets were created as described earlier using terminal spacer fragments carrying ID sequences (SF6-ID-1, SF6-ID-2, SF6-ID-3) unique to the scaffold sets. DC library was produced by folding reaction containing 20 nM scaffold library, DC staple set (100nM of each core staples and C-caps, 500 nM of each biotin staples), 300 nM of ID1 linker (TAAATGGAGGTGAGTGGGAGTAGGGACGAGAACAGGAAAAAAAAAAAAAA) and one-tenth of the final reaction volume of folding buffer (100 mM MgCl2, 50 mM TRIS, 10 mM EDTA) and the overnight temperature ramp described earlier. The folded DC library was purified using three rounds of PEG precipitation and concentration was measured using the Nanodrop 2000 instrument (Thermo Scientific). Oligo(dT)25 Dynabead solution (Thermo Scientific, 61005) with volume equal to the DC sample was washed with storage buffer on a magnetic stand and was subsequently resuspended in the DC sample solution and was incubated on a rotator at room temperature overnight. Beads were pelleted on a magnetic stand and were washed six times in storage buffer after the solution with the unbound DCs were removed. The bound DCs were eluted using toehold mediated strand displacement (TMSD) by incubating the beads in elution solution containing 1.2 μM invader strand (ID1-invader: TTTTTTTTTTTTTTCCTGTTCTCGTCCCTACTCCCACTCA CCTCCATTTA) in storage buffer for 2 hours at room temperature. Finally, the excess invader strand was removed from the eluted DC solution using three rounds of PEG precipitation.

### Preparing stochastic DC library

#### Stochastic insert library generation

An SF6 oligonucleotide carrying a restriction digestion site instead of the ID sequence and a different set of EONs, containing only a single pair of Exchange PAINT docking sites, were used for constructing the stochastic insert library. First spacer libraries were generated from each SF in parallel PCR reactions containing 15 nM of the double stranded SF, 1 μM of forward and reverse primer mixtures consisting of EONs flanking the SF, 200 μM dNTP, 1x GC Buffer and Phusion High-Fidelity DNA Polymerase (New England Biolabs, M0530L) using 30 cycles of PCR. The spacer libraries were purified with agarose gel purification using the MinElute Gel Extraction Kit (Qiagen, 28604). The concentration of the double stranded spacer library stocks was determined using a Nanodrop 2000 instrument (Thermo Scientific). The stochastic insert library was produced by an assembly PCR reaction containing 15nM of each spacer library, 200 μM dNTP, 1x GC Buffer and Phusion High-Fidelity DNA Polymerase (New England Biolabs, M0530L) using 30 cycles of PCR. The assembled insert library was amplified using a PCR reaction containing 1 μM of primers (stochastic_insert_FW:GGAAACAGCTATG ACCATGATTACGAATTCTTGACAGCTAGCTCAGTCCTAGGTATAATG,stochastic_insert_REV:CCCAAGCTTT GTCATGATAATAATGGTTTCTTAGATTTG), 200 μM dNTP, 1x GC Buffer and Phusion High-Fidelity DNA Polymerase (New England Biolabs, M0530L) using 30 cycles of PCR. The insert library was then purified with agarose gel purification using the MinElute Gel Extraction Kit (Qiagen, 28604), and its concentration was measured using a Nanodrop 2000 instrument (Thermo Scientific). The purified insert library was then digested using HindIII HF (New England Biolabs, R3104S) for 30 min at 37 °C followed by heat inactivation at 80 °C for 20 min. The digested insert library was then purified using 1.8x sample volume of AMPure XP beads (Beckman Coulter, A63881) following the manufacturer’s instructions. The random ID sequence was added to the insert library in a ligation reaction containing 10 nM of digested insert library, 0.5µM random ID oligonucleotide (ID random: AGCTNNNNNNNNNNNNNNNNNNNNNNNNNGGATCCTCTAGAGTCGACCTGC) and T4 DNA ligase (New England Biolabs, M0202T) carried out with an incubation at 4°C for 18h followed by heat inactivation step at 65 °C for 10 min. The excess of ID molecules was then removed through a purification step using 1.8x sample volume of AMPure XP beads (Beckman Coulter, A63881) following the manufacturer’s instructions. The concentration of the resulting ID-tagged insert library was measured using Nanodrop 2000 instrument (Thermo Scientific).

#### Stochastic scaffold library generation

The stochastic insert library was cloned into the linearized M13mp18 vector using the CPEC method as described earlier and 2 μL of the cloning reaction was transformed into MegaX DH10B T1R cells by electroporation. The transformation culture after recovery was plated on LB agar plates containing 50 µg/mL Kanamycin and were grown for 48h at 37 °C. The colonies were then recovered from the plates and transferred into 500 mL of growth media (500 mL of 2x Yeast extract Trypton, 1 mL of 50 mg/mL Kanamycin, 4.8 mL of 1M MgCl2, 100 μL of AntiFoam 204) in Thomson’s Ultra Yield™ Flasks and was grown for 18h at 37 °C. The scaffold library was extracted and purified from the produced phage as described earlier.

#### Next generation sequencing of the stochastic scaffold library

1 μg of the scaffold library was converted into double stranded DNA by a Polymerase extension reaction containing 1.2 µM primer (DOCS_random_ONT_primer: CATTATACCTAGGACTGAGCTAGCTGTCAAGAATTCGTAATCATGGTCATAGCTGTTTCC), 1x GC Buffer, 200 µM dNTP and Phusion High-Fidelity DNA Polymerase (New England Biolabs, M0530L) with a 15 min extension step. The double stranded scaffold library was then purified using the NucleoSpin Gel and PCR Clean-up kit (MACHEREY-NAGEL, 740609.50M) following the manufacturer’s instructions. The cleaned double stranded scaffold library was linearized by restriction digestion using EcoRI HF (New England Biolabs, R3101S) followed by a final purification using the NucleoSpin Gel and PCR Clean-up kit (MACHEREY-NAGEL, 740609.50M) following the manufacturer’s instructions.

The library was subsequently prepared for Oxford Nanopore DNA sequencing. The double-stranded DNA was prepared using the manufacturer’s instruction for the Oxford Nanopore Ligation Sequencing Kit 114 (Oxford Nanopore Technologies, SQK-LSK114). Briefly, the DNA was subjected to end repair using the NEBNext FFPE DNA Repair mix (New England Biolabs, M6630S) and NEBNext® Ultra™ II End Repair/dA-Tailing Module (New England Biolabs, E7546S) for 20 °C for 5 min and 65 °C for 5 min. The mix was then added to the ligation buffer and adapter from the LSK-114 kit, supplemented with NEB T4 Ligase (New England Biolabs, M0202S) for 10 min at Room temperature. Following the ligation, fragments were purified using the supplied AMPure beads and the supplied Short Fragment Buffer. The eluted library was then loaded on a MinION Mk1B sequencing device with a R10.4.1 MinION flow cell (Oxford Nanopore Technologies, FLO-MIN114) following manufacturer’s instructions. Base calling was performed within the MinKnow software (v 22.08.4) set to “super accurate base calling”.

#### Challenge-Response experiment with the stochastic DC library

The dominant, unique ID-sequence associated to each unique DC probe was identified from the Nanopore sequencing data of the scaffold and an ID-linker sequence was created for DC(11111) to be used for the earlier described bead-based capture and subsequent TMSD-based release. Folding and capture were performed concurrently using the solid phase synthesis approach for origami structures^50^ using folding reactions containing 100nM DP library, 270mM Sodium polytungstate, DC staple set (100nM of each core staples and C-caps, 500nM of each biotin staples) and 50nM ID-linker (ID(11111)_linker, GGGCAATTGTGACATTGATAAACTAGTGCGTCTCCCAAAAAAAAAAAAAA). The folding reaction was mixed with 0.5 reaction volume of pelleted Oligo(dT)25 Dynabead solution (Thermo Scientific, 61005) and was carried out with the overnight temperature ramp described earlier. Beads were washed and bound DCs were eluted using 1.2μM invader strand (ID(11111)_invader, TTTTTTTTTTTTTTGGGAGACGCACTAGTTTATCAATGTCACAATTGCCC) as described earlier.

### Data encoding and decoding

Given the number of colors, we calculated the number of color combinations n as 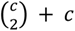. For Huffman encoding, we used a publicly available implementation (obtained from https://github.com/rdbliss/huffman) to build n-ary Huffman trees using the frequencies of letters in English texts (obtained from: https://norvig.com/mayzner.html). Each message was cleaned up to remove all but the 26 letters in the English alphabet, which were all converted to lowercase.

For zlib encoding, we converted the message to bytes using utf-8 encoding, and compressed these using the zlib library in python at compression level 9. The bytes were then converted to an integer and converted to base n using gmpy2^51^.

The encoded message was then divided into overlapping k-mers, where each k-mer would be held by one data carrier. The overlap of each sequential k-mer was set to k-s, where s is the shift. The shift was 1 for all experiments except the encoding of the “alice” text, where it ranged from 1 to 4. If the final k-mer was too short, it was filled to the required length with a predetermined filler character (“0”).

Decoding obtained messages (see below on how they were obtained) was done using the inverse operation. Zlib-encoded messages were converted to an integer, which was then converted to bytes and decompressed. For shifts larger than 1, if zlib-decoding failed and the decoded message ended with the filler character, it would be retried after removing the filler character. This would be repeated until decoding was successful, or until the message no longer ended with the filler character. Huffman-encoded messages were directly decoded using the same Huffman tree used for encoding.

### Structure count simulations

Relative observation counts for each structure were drawn from a power-law distribution obtained from fitting experimental data using the stats module from scipy. Relative observation counts were then adjusted upwards into real observation counts so that the lowest observed structure would have 100 observations. For physically corrected errors, which had a consistent error rate, we precalculated the fraction of structure derivatives differing by 1 to k characters. Then, for each structure, the number of derivatives was randomly drawn from a Poisson distribution with [lambda = count * fraction_errors], and that many derivatives were generated and added to the observations. The same number of observations were subtracted from the unaltered observations. For uncorrected errors, which had a variable error rate, we recursively generated all structure derivatives and their relative observation rates. When the relative rate would drop below a cutoff (1e-15), that structure and its derivatives would no longer be considered. We then used a Poisson distribution to draw real observations from the relative observation rates.

### Algorithmic error correction

Algorithmic error correction was inspired by earlier described sequence error correction strategies (UMI-tools)^52^. First, a directed graph is created from the structures, where an edge is formed from structure a to b if structure a and b differ at most one character, and if a has a higher observed count than b. A depth-first search is then performed on the graph to find all connected groups, starting from the most abundant structures. The most abundant structure per group is considered the correct structure, while the others are removed.

### Message extraction

Starting from structure abundance data, optionally after algorithmic error correction, a weighted directional graph was created. Given a shift parameter s, each structure was split into two nodes, a left node (containing all but the last s characters) and a right node (containing all but the first s characters). An edge with weight equal to the structure abundance was added connecting the left node to the right node.

Putative start nodes were identified as those nodes with no incoming edges. If no such nodes were present, all junction nodes (defined as nodes with at least one incoming edge and two outgoing edges, or vice versa) were considered start nodes. The graph was then compressed by keeping only junction nodes, start nodes and end nodes and compressing all intermediate connections between them into a single edge each.

Starting from the putative start nodes, paths in the collapsed graph were searched recursively. During this search, paths were scored by accumulating the observed abundance of each structure that is part of that path. The score per structure was multiplied by a penalty factor when selecting structures multiple times. This factor scaled linearly from 0.1 to 0.9 with the logarithm of the structure’s abundance. If the structure was selected more than twice, the factor was raised to the power of the number of times the structure was selected, minus one.

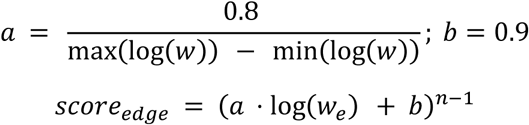

Where w_e_ is the weight of an edge (structure abundance) and n is the number of times that edge is used in the path. If extending a path resulted in a decrease in overall score twice in a row, the path was pruned. This strategy allowed paths to reselect multiple nodes twice, if, after that, extending the path would result in a score increase (by selecting previously unselected nodes). Path finding for a given start node was aborted if the length of the path in the collapsed graph was more than 1000 nodes (enforced by a recursion limit). Path finding and decoding were limited to 10 minutes each, keeping only the results obtained within that timeframe.

To calculate the relative probability of each path, we assumed that real observations and erroneous observations both were distributed as Normal distributions with the same standard deviation, but different mean values:

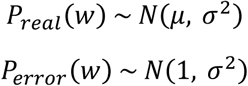

Where µ and sigma were calculated from the input data. For each edge, we then calculated the normalized probability of it belonging to either distribution.

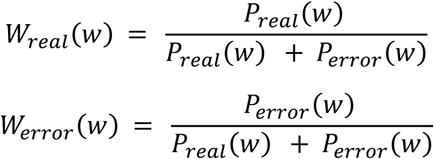

For each path p, the total likelihood of that path was calculated by multiplying W_real_ for all edges that were part of the path and multiplying W_error_ for all edges not part of the path.

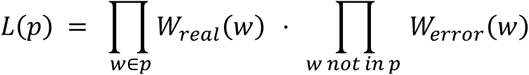

All total likelihoods were then normalized, either by all paths, or by all paths in the same connected group.

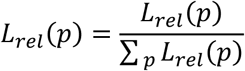

### NGS sequencing of the inserts and scaffolds

The sequencing of the DOCS insert was performed with an ONT MinION Mk1B using a FLO-MIN114 flow cell. Sequencing library was prepared from a purified insert using the ligation sequencing DNA V14 kit (ONT, SQK-LSK114) and following the manufacturer’s instructions.

### Sequence analysis of the inserts

Nanopore data of the DOCS insert was processed with an in-house script. Sequences sharing terminal 8bp with the reference sequence were extracted from the reads and those with selected size distribution corresponding to the main observed products were grouped. Sequences in respective groups were aligned using CLUSTAL 2^53^ and a consensus sequence was generated from them.

### Sequence analysis of random DOCS library

Nanopore reads were aligned using minimap2^54^ with the “map-ont” preset to a reference insert where the IDs and color sequences were replaced with Ns. The regions corresponding to the IDs and color sequences were extracted with in-house scripts. Each color sequence was aligned to each of the three reference color sequences using Biopython’s PairwiseAligner^55^, with the following settings: mode set to local, match score set to 2.0, mismatch score set to –3.0, gap score set to 0.0, internal gap open score set to –5.0, internal gap extend score set to –2.0. A color sequence was assigned to the reference color sequence it had the highest alignment score with, if a) the score was at least 0.3 times the maximum score, and b) the highest score occurred only for one color sequence.

## Acknowledgements

This work was supported by the European Union’s EIC Pathfinder Challenges 2022 programme (DuraStore – 101115410).

## Author contributions

F.F., A.K. and B.H. conceived the project. F.F., A.K. and A.L. designed the experiments. A.K. developed the data encoding, decoding and simulation pipeline. F.F., A.K., A.L., B.S. and I.B. carried out the experiments. F.F., A.K. and A.L. analyzed the data. F.F., A.K., A.L., B.S., I.B. and B.H. wrote the manuscript. All authors discussed the results and commented on the manuscript.

## Use of AI declaration

An LLM was used to assist in editing the text for clarity and conciseness. The authors reviewed and take full responsibility for all content.

