## Supplementary figures for "A combinatorial DNA origami platform for biologically replicable, thermostable data storage and molecular authentication"

Supplementary materials

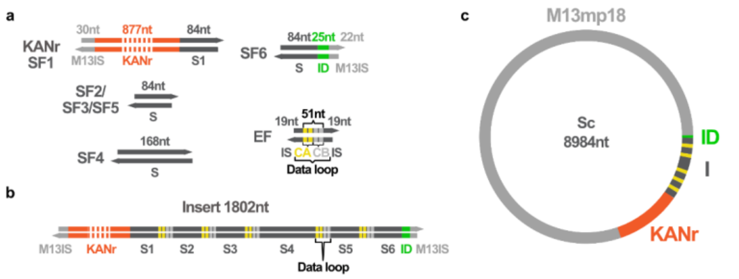


***Supplementary Fig. 1. Size and composition of DOCS building blocks, the insert and the scaffold constructed from them a,*** *Inserts are constructed from two types of building blocks spacer fragments (SF) and data encoder fragments (DEF). SFs contain spacer sequences (S) of equal size, except SF4 to introduce asymmetry aiding the decoding process. Terminal SFs contain an M13 insertion sequence (M13IS), complementary to the vector needed for the cloning and SF6 contains the ID sequence (ID) used for random access. DEFs consist of data encoding sequences (DES) that contain pairs of Exchange PAINT docking sites (channel A (CA) and a "blank" (CB) docking sites in this example) flanked by insertion sequences (IS) complementary to adjacent SFs in the insert.* ***b,*** *The strand architecture of the fully assembled insert and its size.* ***c,*** *The map of the assembled scaffold (Sc) and its size.*

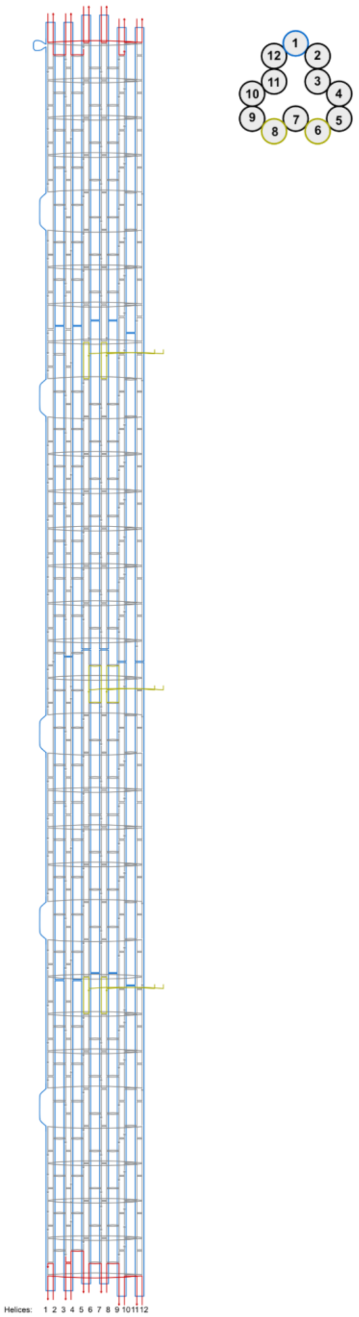


***Supplementary Fig.2 Design schematic of the data carrier origami structure*** *Strand diagram of the data carrier with helices numbered, showing the core (grey), A or C-caps (red) and biotin anchor (yellow) staple strands that fold the custom scaffold molecule (blue) into a 12-helix bundle origami structure leaving the data encoding sequences (DES) and the ID sequence as unpaired regions. (center) Cross section of the data carrier with helices represented as circles following the same helix numbering convention as the strands diagram with helices containing biotinylated protrusions highlighted with yellow and the helix with the DES and ID sequences highlighted with blue).*

GGAAACAGCTATGACCATGATTACGAATTCTTGACAGCTAGCTCAGTCCTAGGTATAATGCTAGCTACTAGAGAAAGAGGAGAAATACTAGATGAGCCAT

ATTCAACGGGAAACGTCTTGCTCCAGGCCGCGATTAAATTCCAACATGGATGCTGATTTATATGGGTATAAATGGGCTCGCGATAATGTCGGGCAATCA

GGTGCGACAATCTATCGATTGTATGGGAAGCCCGATGCGCCAGAGTTGTTTCTGAAACATGGCAAAGGTAGCGTTGCCAATGATGTTACAGATGAGAT

GGTCAGACTAAACTGGCTGACGGAATTTATGCCTCTTCCGACCATCAAGCATTTTATCCGTACTCCTGATGATGCATGGTTACTCACCACTGCGATCCC

CGGGAAAACAGCATTCCAGGTATTAGAAGAATATCCTGATTCAGGTGAAAATATTGTTGATGCGCTGGCAGTGTTCCTGCGCCGGTTGCATTCGATTCC

TGTTTGTAATTGTCCTTTTAACAGCGATCGCGTATTTCGTCTCGCTCAGGCGCAATCACGAATGAATAACGGTTTGGTTGATGCGAGTGATTTTGATGAC

GAGCGTAATGGCTGGCCTGTTGAACAAGTCTGGAAAGAAATGCATAAGCTTTTGCCATTCTCACCGGATTCAGTCGTCACTCATGGTGATTTCTCACTT

GATAACCTTATTTTTGACGAGGGGAAATTAATAGGTTGTATTGATGTTGGACGAGTCGGAATCGCAGACCGATACCAGGATCTTGCCATCCTATGGAAC

TGCCTCGGTGAGTTTTCTCCTTCATTACAGAAACGGCTTTTTCAAAAATATGGTATTGATAATCCTGATATGAATAAATTGCAGTTTCATTTGATGCTCGAT

GAGTTTTTCTAAATAAACCAGCCAGCCGGAAGGGCCGAGCGCAGAAGTGGTCCTGCAACTTTATCCGCCTCTTCGCCAGTTAATAGTTTGCGCAACTT

TTCTTCATTATTTTCTTCATTATTTTCAATGTATTTTTCAATGTATTTTGAAAACTCACGTTAAGGGATTTTGGCCAAGTCATTCTGAGAATAGTGTATGCGG

CGACCGGGCTTACCATCTGGCCCCAGTGCTTTTTCAATGTATTTTTCAATGTATTTTCTACAATAATTTCTACAATAATTTTCATTCAGCTCCGGTTCCCAA

CGATCAAGGCGAGTTACATGATCCCCCATGTTGTGCAATAAGTTGGCCGCAGTGTTATCACTCTTTTCTTCATTATTTTCTTCATTATTTCTACAATAATTT

CTACAATAATTTTTTTCTGTGACTGGTGAGTACTCAATCATGAGATTATCAAAAAGGATCTTCACCTAGATCCTTTTAAATTAAAAATGAAGTTTTAAATCAA

TCTAAAGTATATATGAGTAAACTTGGTCTGACAGTTACCAATGCTTAATCAGACATAGCAGAACTTTAAAAGTGCTCTTTATCTACATATTTATCTACATATT

TCTACAATAATTTCTACAATAATTTAGATCCAGTTCGATGTAACCCACTCGTGCACCCAACTGATCTTCAGCATCTTTTACTTTCGCAAAAAAGGGAATAA

GGGCGACATTTATCTACATATTTATCTACATATTTTCAATGTATTTTTCAATGTATTTTCAGGGTTATTGTCTCATGAGCGGATACATATTTGAATGTATTTAG

AAAAATAAACAAATCTAAGAAACCATTATTATCATGACAAAGCTCCCTACTCCCACTCACCTCCATTTAGGATCCTCTAGAGTCGACCTGC

***Supplementary Fig 3. Consensus sequence of insert (52104) product and side product*** *Reference sequence of insert(52104) insert with the resistance cassette sequence highlighted in green and the sequence start sites for the product (black) and side product (grey) highlighted.*

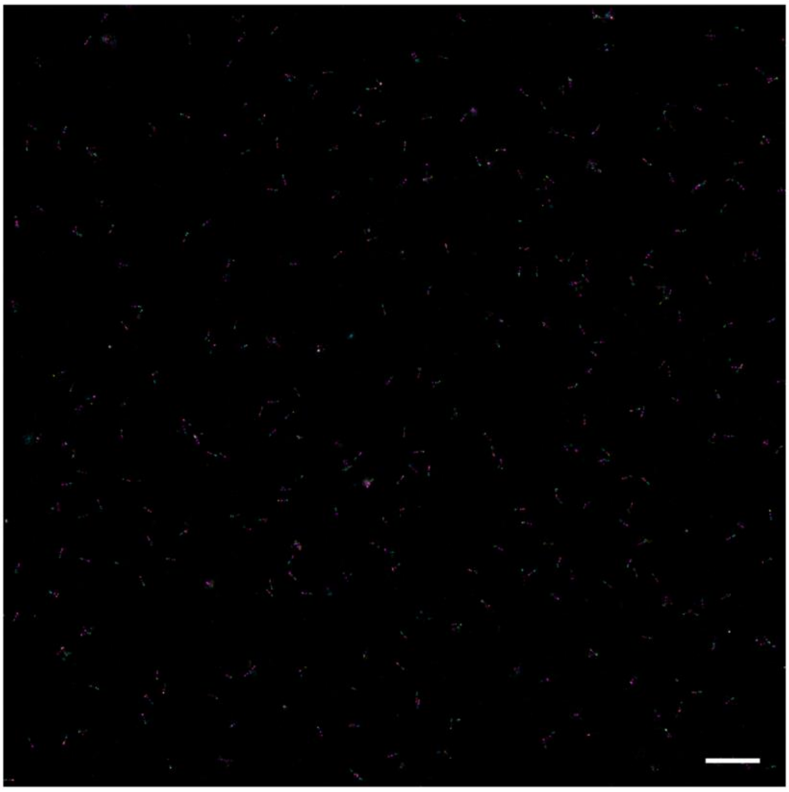

***Supplementary Fig 4. Field of view Exchange PAINT image of senary data encoding data carriers (scale bar = 1µm)***

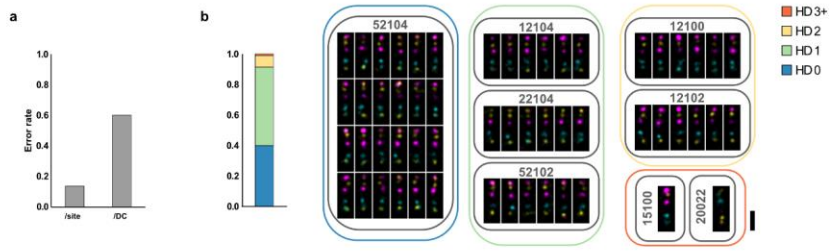


***Supplementary Fig 5. Error rates and mutational analysis of data carrier (DC) encoding senary information a,*** *Bar plot showing the error rates per DC site (0.14) or DC (0.60) extracted calculated from the processed Exchange PAINT data* ***b,*** *Bar plot showing the Hamming distance (HD) distribution of the decoded DCs compared to the encoded senary string (52104) showing the relative abundance of DCs with codes of given HD from the reference (HD0: 0.399, HD1: 0.514, HD2: 0.077, HD3+:0.009). (left) DNA PAINT images of decoded DCs with codes in indicated HD from the reference string (52104) (scale bar = 100nm). (right)*

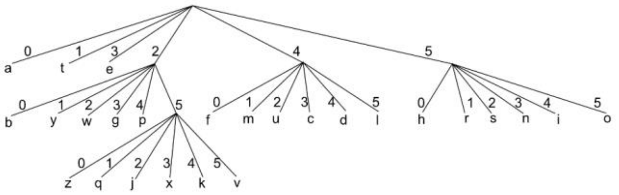


***Supplementary Fig 6. Huffman-tree used for the encoding of text information into senary format.***

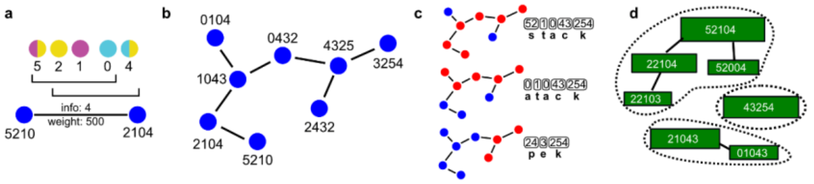


***Supplementary Fig 7. Data reconstruction and algorithmic error correction. a,*** *Each observed structure is converted to two nodes formed from the left and right flanks, connected by an edge weighted by the structure abundance.* ***b,*** *Converting all structures to nodes forms a directed graph.* ***c,*** *A path-finding algorithm traverses all paths and scores them by accumulated abundance accounted for. Each message is then decoded.* ***d,*** *Optionally, before graph creation, structures are fused together if their sequences lie within one hamming distance. The most abundant in the group is assumed to be the correct structure.*

**Commented [FF1]:** Change structure to DC?

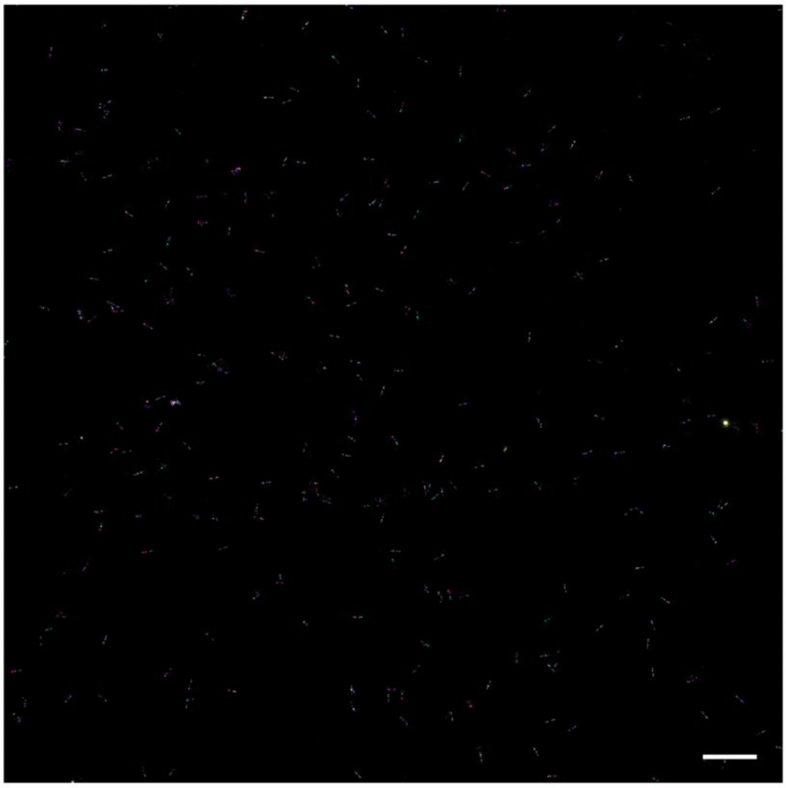

***Supplementary Fig 8. Field of view Exchange PAINT image of text encoding DC library (scale bar = 1µm)***

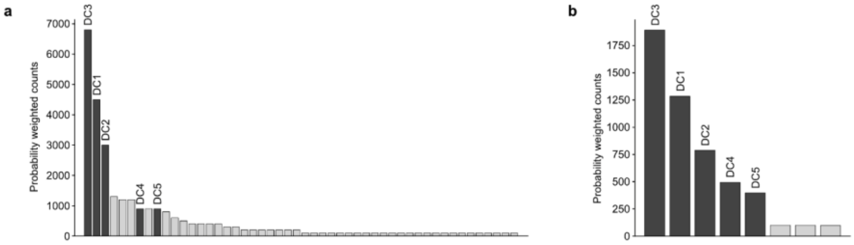


***Supplementary Fig 9. Frequency distribution of DCs in the text encoding library*** *Plots showing the probability weighted counts of DCs encoding the text "stack" (DC1-5) in dark gray and DCs with unexpected senary strings generated by errors in light gray, detected in the Exchange PAINT data through processing (**a,**) without error correction and (**b,**) with physical error correction (PEC).*

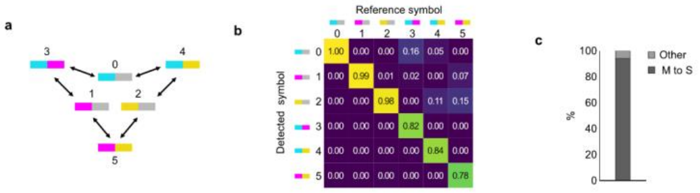


***Supplementary Fig 10. Error analysis of text encoding DC library and physical error correction (PEC)** **a,** The dominant error pathways if only channel dropouts are permitted. **b**, Error rate matrix showing the rate of symbols detected compared to the intended symbols displaying a high rate of correctly identified symbols and an elevated error rate for mixed channel symbols (3-5) converting to corresponding single channel symbols (0-2) supporting the hypothesis of channel dropouts being the dominant source of detection error. **c,** Plot displaying percentage-wise contribution of error types (Mixed to single (MS), Other) to all detected errors.*

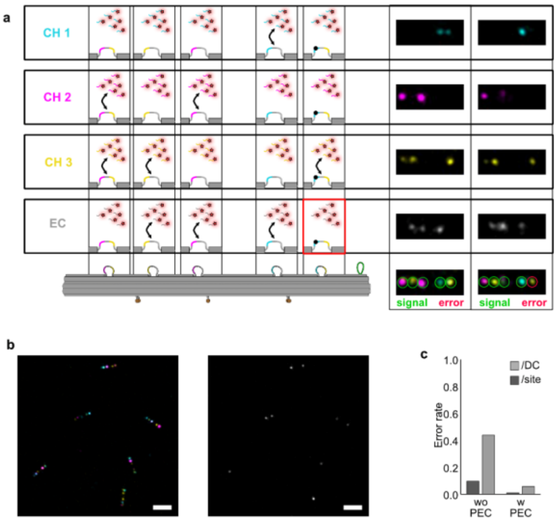


***Supplementary Fig 11. Physical error correction (PEC) strategy** **a,** Physical error correction (PEC) strategy, where a fourth channel is used with imaging sites complementary to the empty placeholder sequence in single channel sites, which helps distinguishing genuine and error generated single channel positions. **b**, Exchange PAINT image of single word ("stack") encoding DC library (left) and its corresponding error correcting channel (right) (scale bars = 200nm) **c**, Plot showing the error rates per DC (light gray) and per DC site (darks grey) for the single word ("stack") encoding DC library with and without utilization of PEC.*

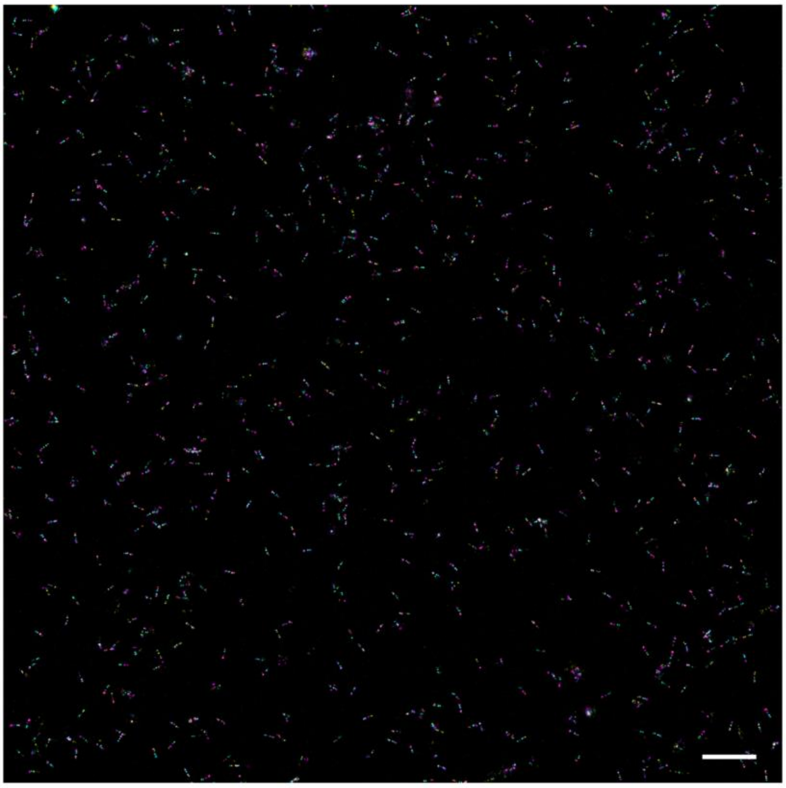

***Supplementary Fig 12. Field of view Exchange PAINT image of text encoding DC library recloned using scaffold library (scale bar = 1µm)***

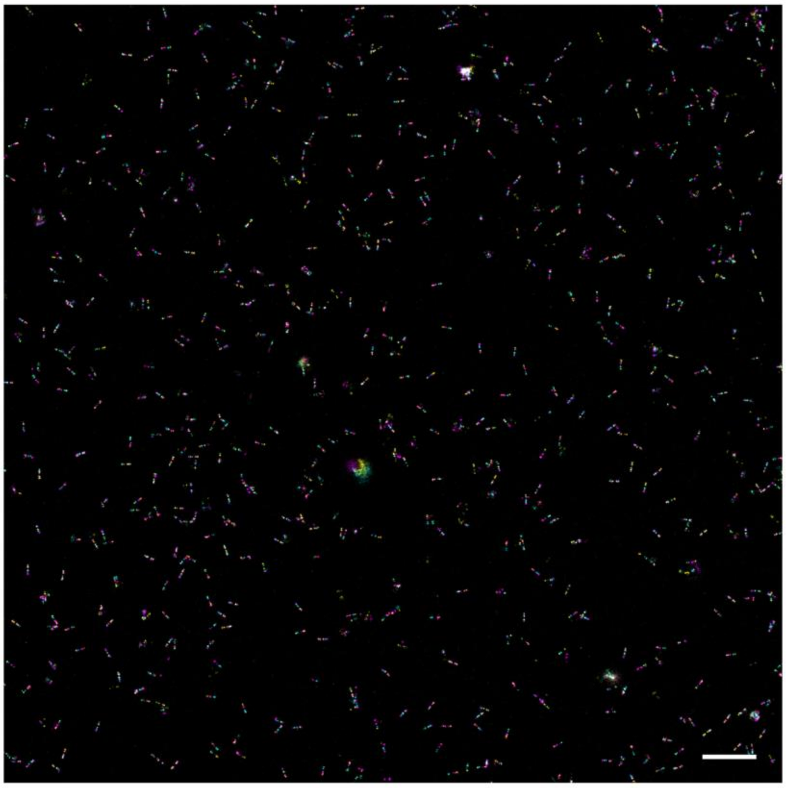

***Supplementary Fig 13. Field of view Exchange PAINT image of text encoding DC library recloned using DC library (scale bar = 1μm)***

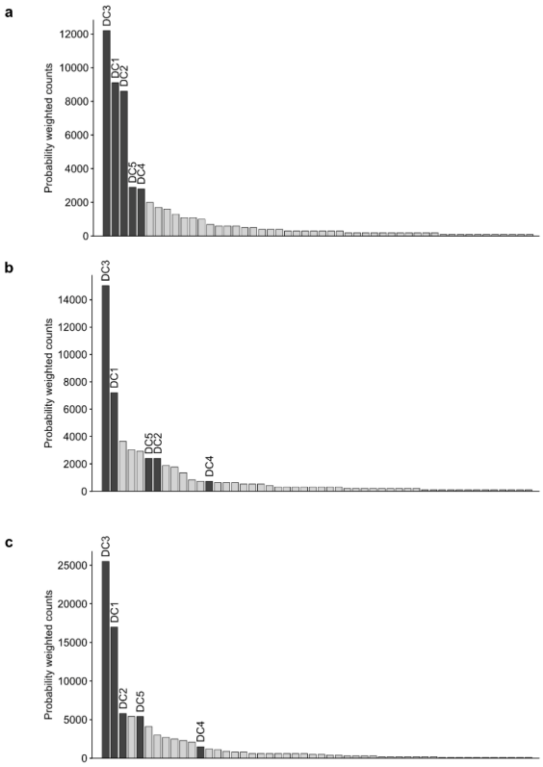


***Supplementary Fig 14. Frequency distribution of DCs in the libraries replicated via molecular cloning*** *Plots showing the DCs encoding the text "stack" (DC1-5) in dark gray and DCs with unexpected senary strings generated by errors in light gray, detected in the Exchange PAINT data recorded of the (**a,**) reference library, (**b,**) the library cloned from the scaffold library and (**c,**) the library cloned from the DC library through processing without error correction.*

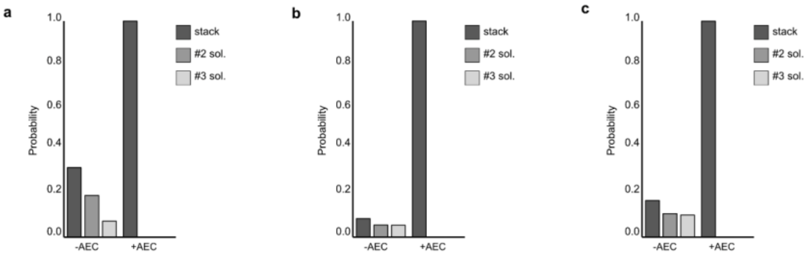


***Supplementary Fig 15. Recovery probabilities of messages encoded in libraries replicated via molecular cloning*** *Plots showing the calculated probability of the top three decoding solutions from the data recovered from the DP library using combinations of the different error correction strategies.*

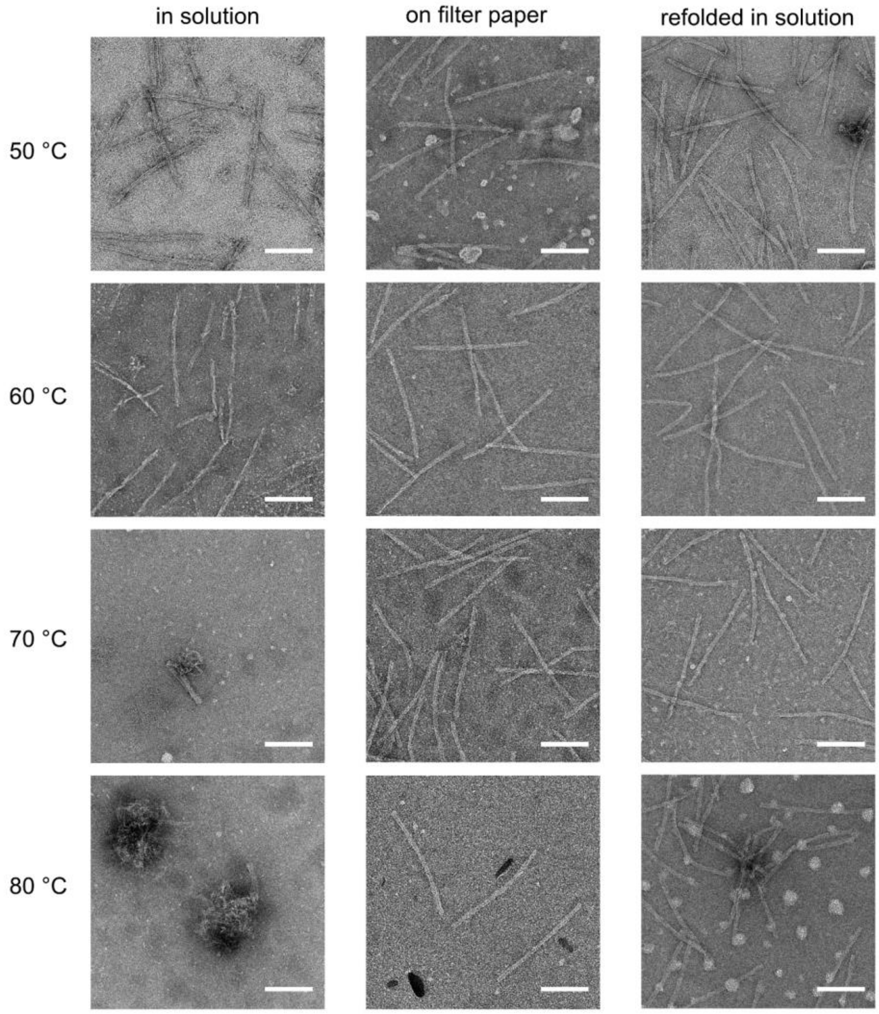

***Supplementary Fig 16.*** **TEM images of the DC library incubated at elevated temperatures** *(scale bars = 100nm)*

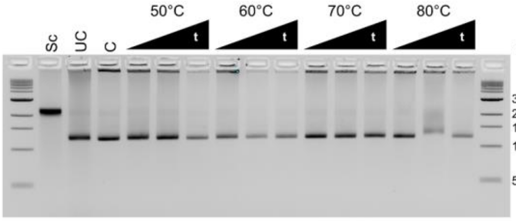


***Supplementary Fig 17. DC library refolded from dried form*** *Agarose gel electrophoresis of DC library (DC1-5) samples incubated at 50-80°C for 15-60 minutes in dried pellet form and refolded afterwards along with scaffold library (Sc), uncleaned (UC) and cleaned (C) control DC library (DC1-5) samples.*

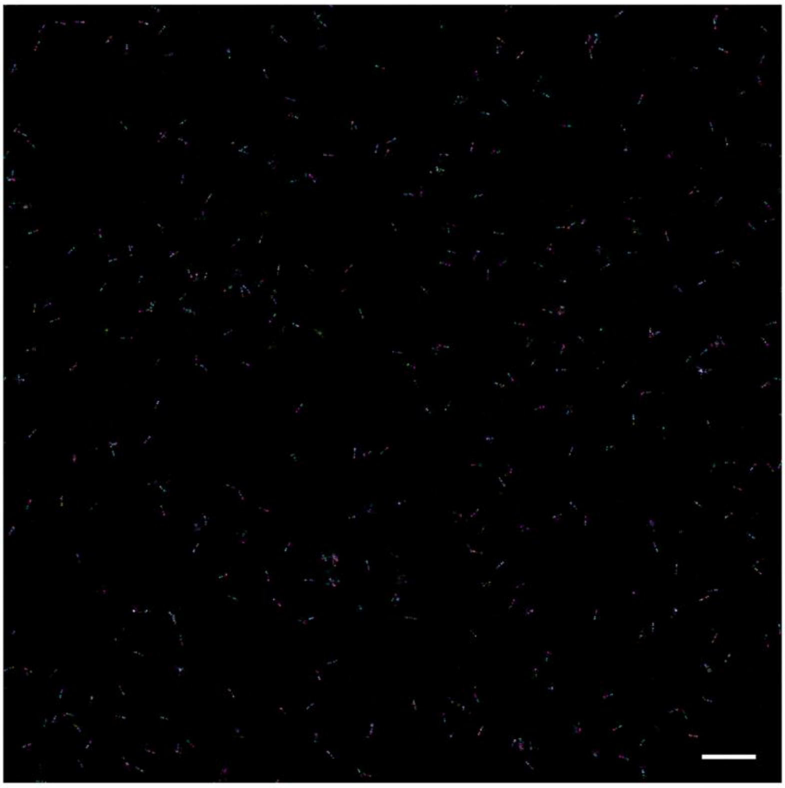

***Supplementary Fig 18. Field of view Exchange PAINT image of text encoding DC library incubated at 80°C for 1h on filter paper (scale bar = 1µm)***

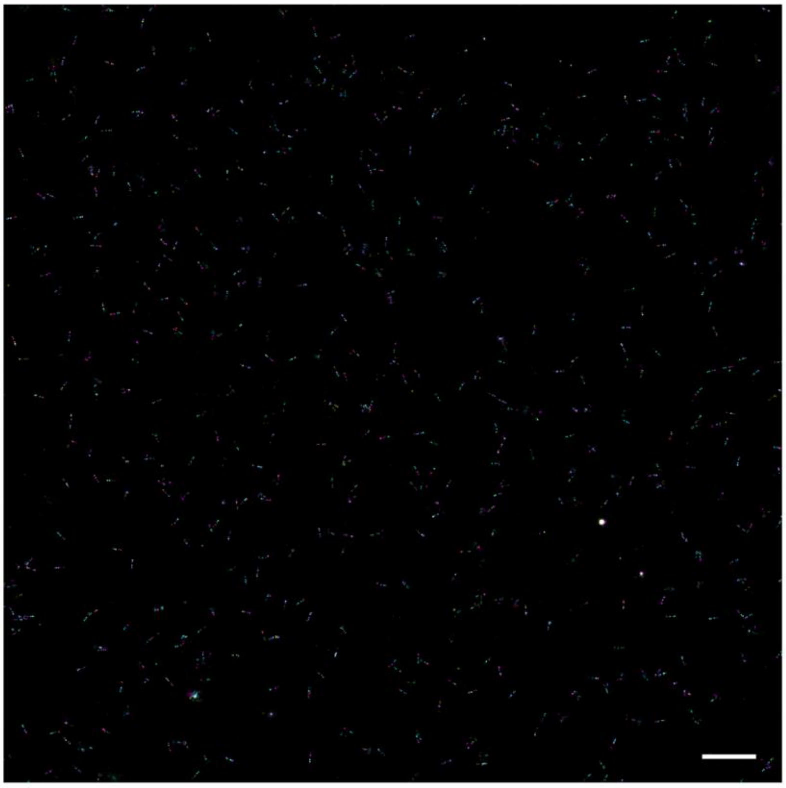

***Supplementary Fig 19. Field of view Exchange PAINT image of text encoding DC library incubated at 80°C for 1h in buffer and subsequently refolded (scale bar = 1μm)***

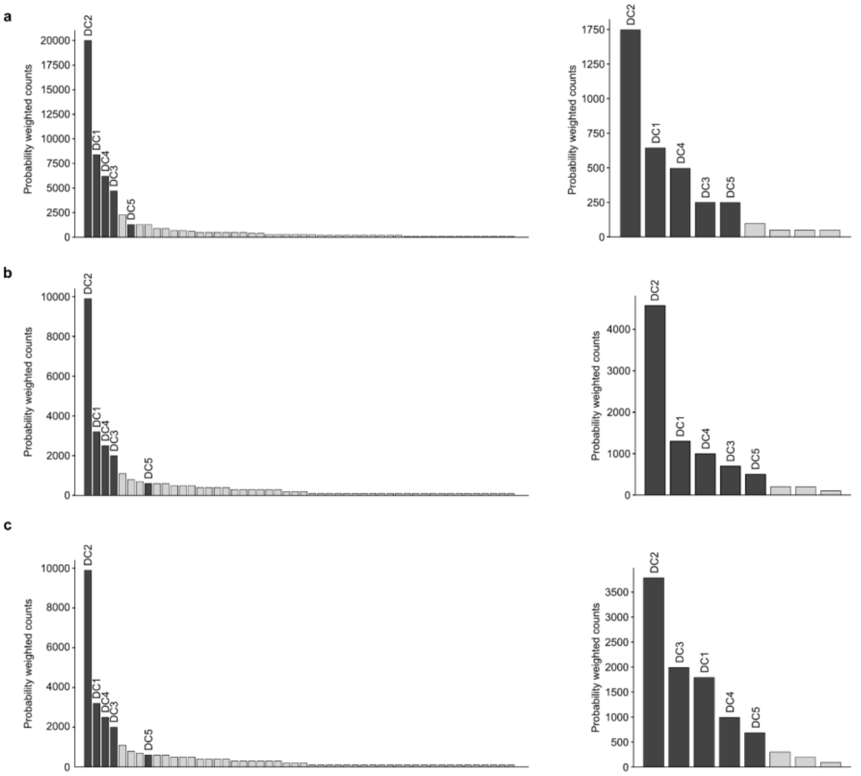


***Supplementary Fig 20. Frequency distribution of DCs incubated at high temperatures*** *Plots showing the DCs encoding the text "stack" (DC1-5) in dark gray and DCs with unexpected senary strings generated by errors in light gray, detected in the Exchange PAINT data recorded of the (****a,****) reference library, (****b,****) the library incubated at 80°C for 1 hour on filter paper and (****c,****) in liquid and refolded afterwards through processing without (left) or with physical error correction (right).*

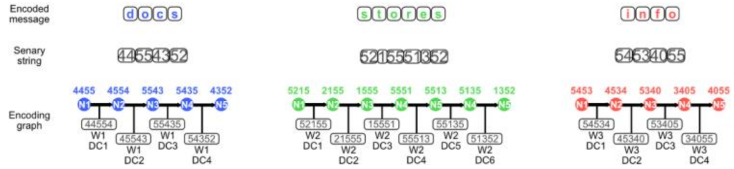


***Supplementary Fig 21. Encoding scheme for 3-word data set*** *Text messages, their senary form and the graph generated from them with their corresponding DCs.*

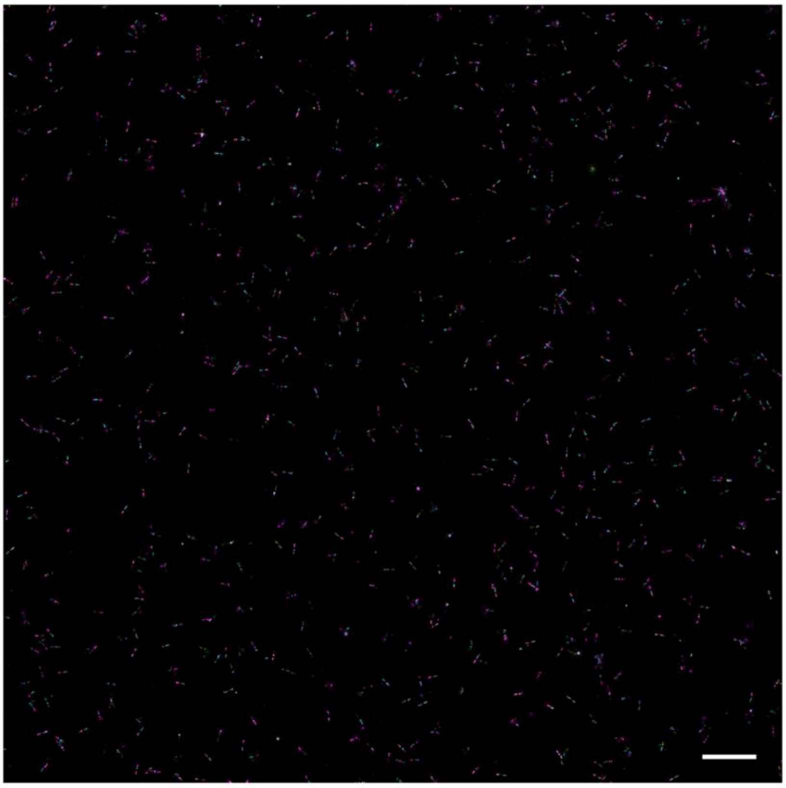

***Supplementary Fig 22. Field of view Exchange PAINT image of multi-file encoding DC library (scale bar = 1µm)***

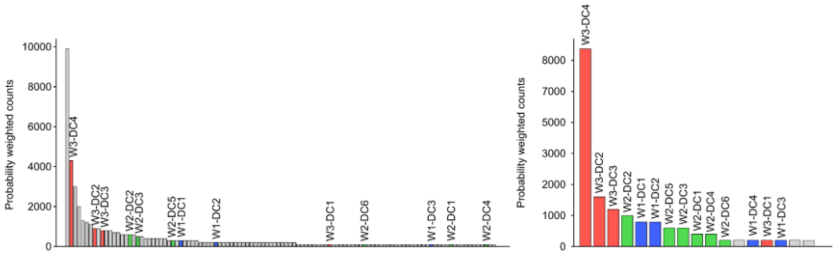


***Supplementary Fig 23. Frequency distribution of multi-file encoding DC library*** *Plots showing the data carriers (DCs) encoding the text "docs" (W1-DC1-4), "stores" (W2-DC1-6) and "info" (W3-DC1-4) in blue, green and coral respectively and probes generated by errors in light gray detected in the Exchange PAINT through processing without (left) or with physical error correction (right).*

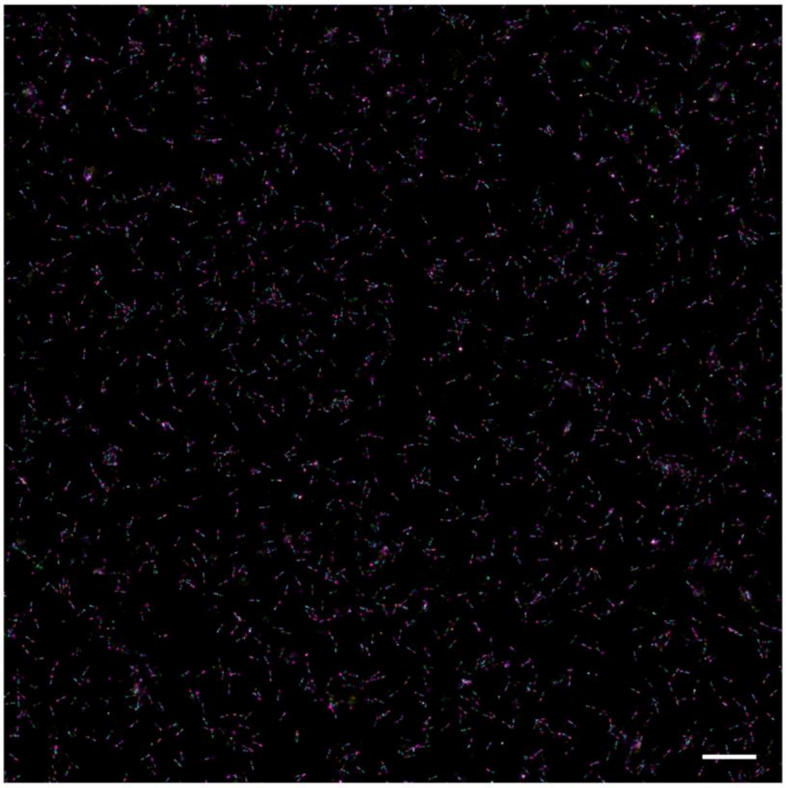

***Supplementary Fig 24. Field of view Exchange PAINT image of multi-file encoding DC library after random-access selection (scale bar = 1µm)***

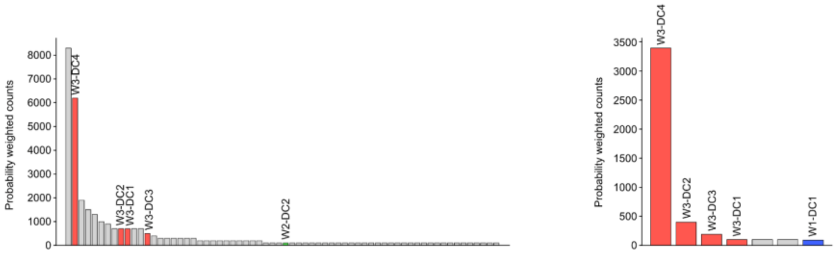


***Supplementary Fig 25. Frequency distribution of multi-file encoding DC library after selection*** *Plots showing the multi-file encoding DC library after selection with DCs encoding the text "docs" (W1-DC1-4), "stores" (W2-DC1-6) and "info" (W3-DC1-4) in blue, green and coral respectively and probes generated by errors in light gray detected in the Exchange PAINT through processing without (left) or with physical error correction (right).*

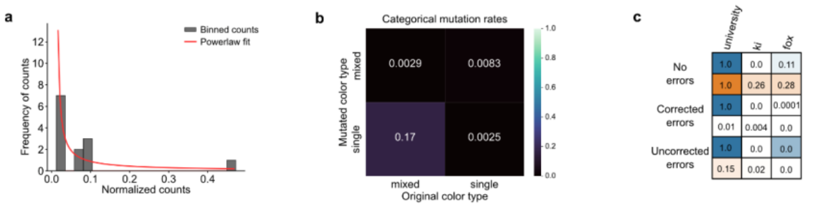


***Supplementary Fig 26. Simulation parameters and text storage without AEC a,*** *Plot showing the DC distribution bias observed in experimental data estimated by a power law function used for the simulations* ***b,*** *Plot showing the error rates extracted from experimental data used for the simulations* ***c,*** *Relative probabilities of correct path over all other paths in graphs obtained from simulations for DOCS encoded texts using Huffman encoding (orange) or zlib encoding (blue).*

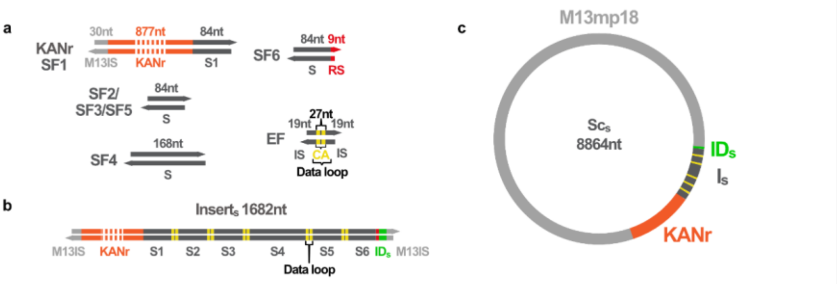


***Supplementary Fig. 27. Size and composition of DOCS building blocks, the insert and the scaffold used for building the stochastic DOCS library*** *a, Stochastic inserts are constructed from similar building blocks as the conventional insert molecules: spacer fragments (SF) and data encoder fragments (DEF). SFs sequences are nearly identical to those used for the conventional DI construction except that SF6 contains a restriction site (RS) for the addition of the random ID molecule at later steps instead of the fixed ID sequence. The DEFs also have the same composition except here a smaller data encoding sequences (DES) was used with a single pair of Exchange PAINT docking sites (channel A (CA) docking sites in this example).*

*b, The strand architecture of the fully assembled, ID-labelled stochastic insert and its size. c, The map of the stochastic scaffold ($Sc_S$) and its size.*

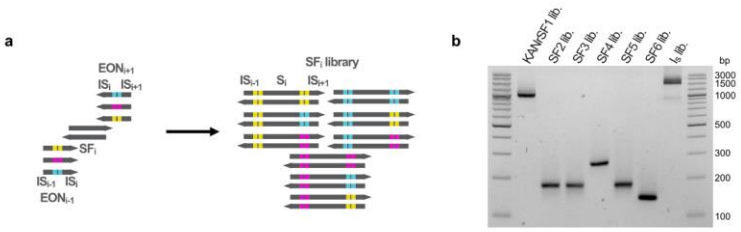


***Supplementary Fig. 28. Construction of the components used for the one-pot construction of the stochastic insert library*** *a, Spacer libraries are created by PCR amplification of a given spacer fragment molecule ($SF_i$) with the mixture of the EONS of the sites flanking the given SF with complementary insertion sequences ($IS_i$). b, Agarose gel electrophoresis of the spacer fragment libraries (KANr-SF1, SF2-SF6) and the stochastic insert constructed from them ($I_S$).*

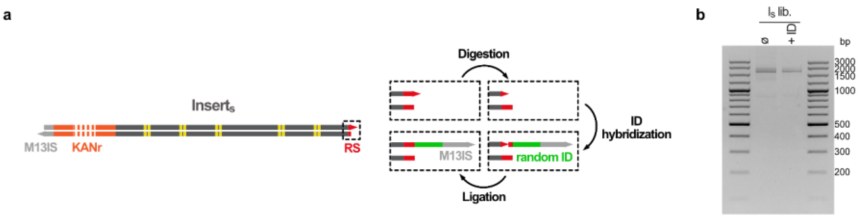


***Supplementary Fig. 29. The tagging of the stochastic insert library with the random ID molecule*** *a, The stochastic insert ($Insert_S$) library contains a terminal restriction site (RS), which is used for the addition of the random ID molecule by subsequent digestion, hybridization and ligation steps. b, Agarose gel electrophoresis of the stochastic insert library ($I_S$) before (ø) and after (+ID) the addition of the random ID molecule.*

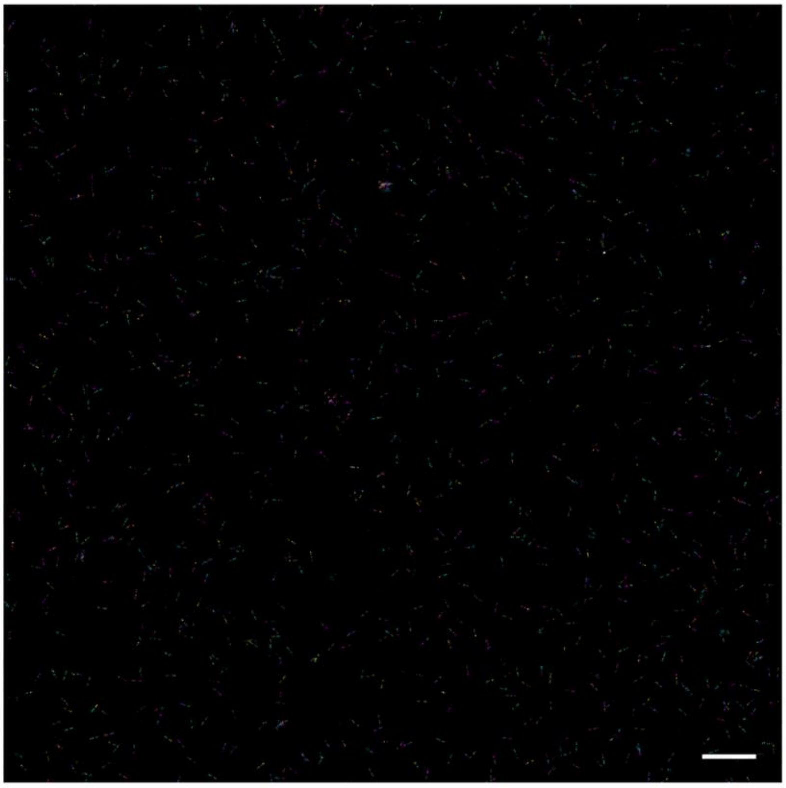

***Supplementary Fig 30. Field of view Exchange PAINT image of stochastic DC library (scale bar = 1µm)***

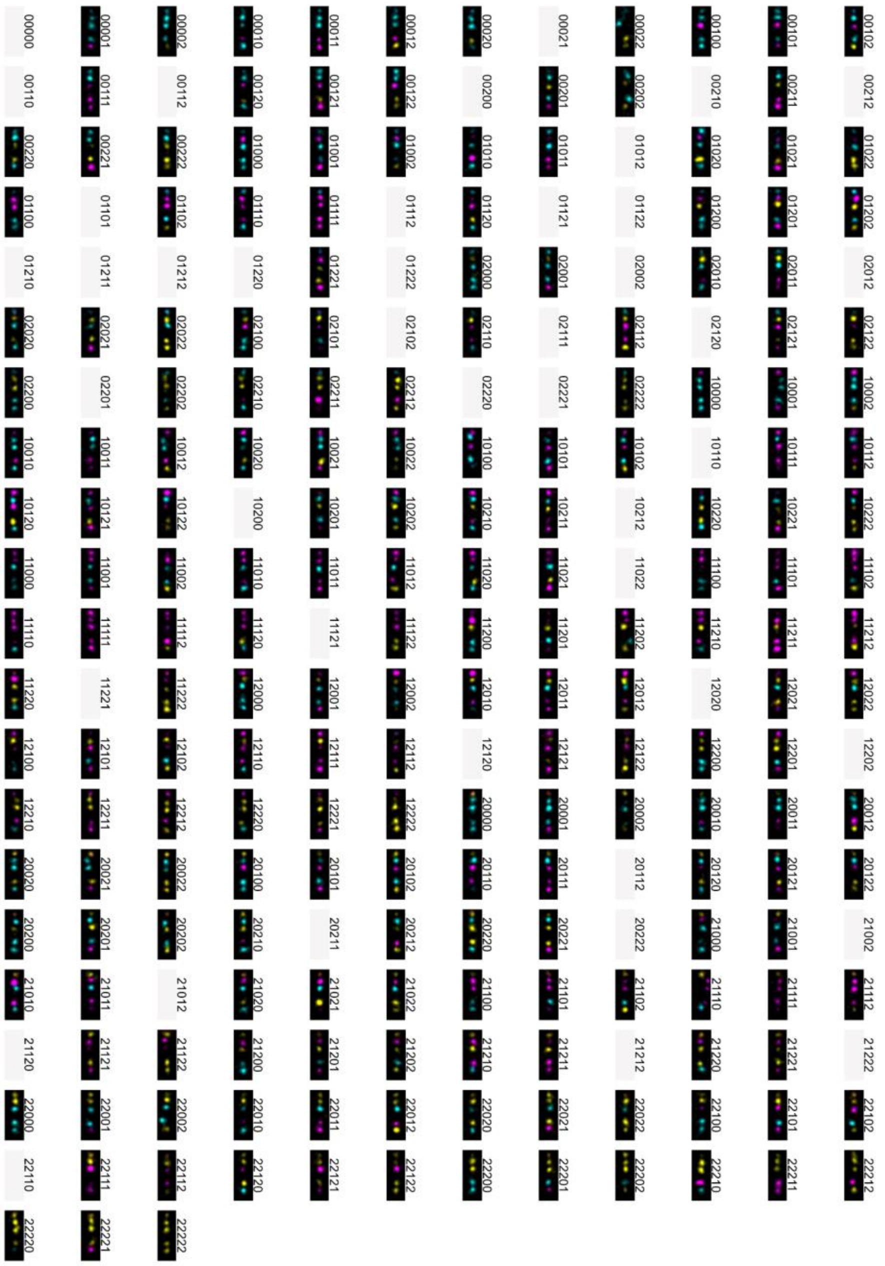

***Supplementary Fig. 31. Individual DCs in the stochastic DC library*** *Exemplary Exchange PAINT images of the unique DCs detected in the PAINT data of the stochastic DC library along with their code with place holders used for DCs not detected in the dataset.*

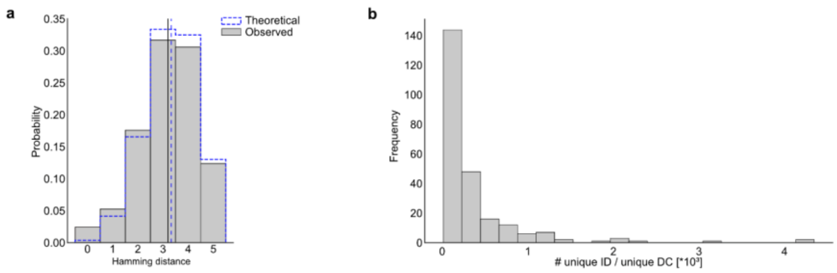


***Supplementary Fig 32. NGS characterization of the stochastic DC library a,*** *Plot showing the distribution of the pairwise Hamming distances of the codes of DCs in the stochastic, 3-color library with the theoretical distributions and its mean shown with blue dashed lines and the frequency weighted distribution of the sample and its mean in grey.* ***b,*** *Plot showing the distribution of the number of unique ID sequences observed per unique DC codes in the library.*

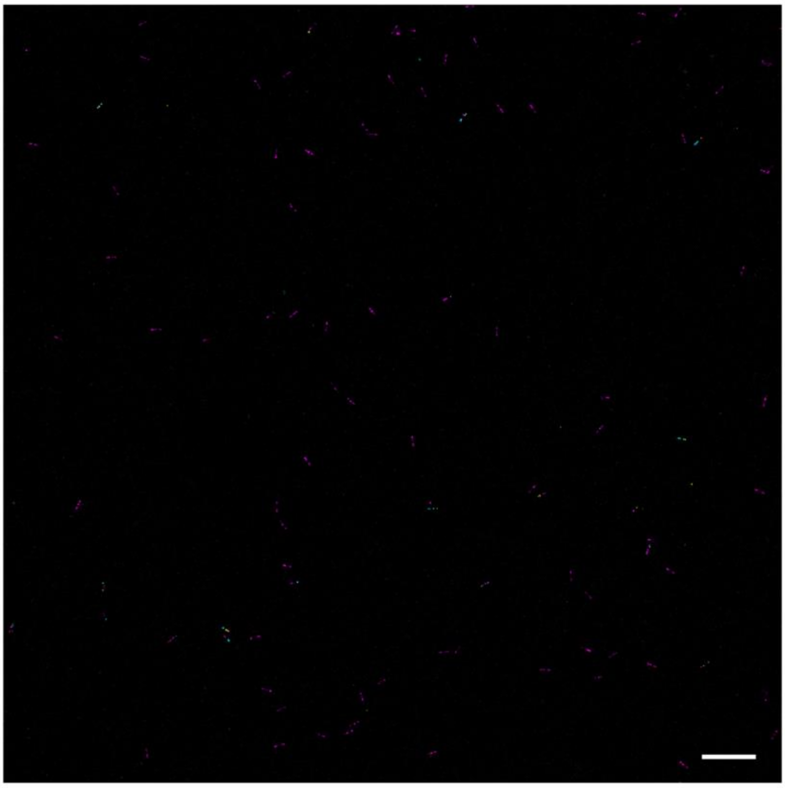

***Supplementary Fig 33. Field of view Exchange PAINT image of the response sample produced by the challenge performed on the stochastic DC library (scale bar = 1µm)***

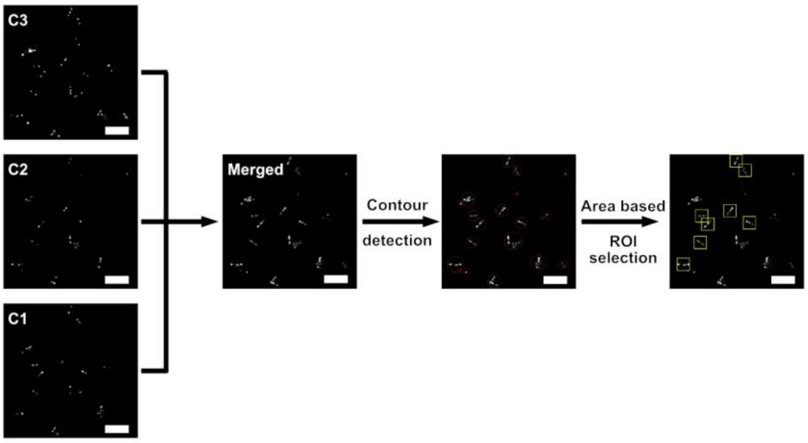


***Supplementary Fig 34. DC detection and data extraction from Exchange PAINT data*** *The workflow of the part of the Exchange PAINT processing pipeline used for detecting individual DCs and extracting events produced by them by contour detection and subsequent are based contour selection in the merged data set (scale bars = 500nm).*

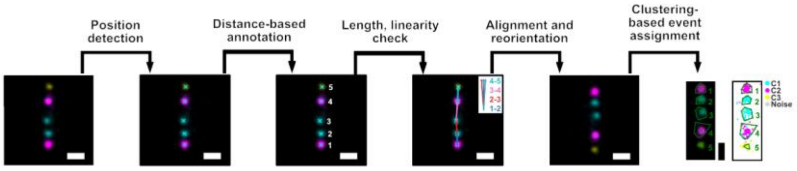


***Supplementary Fig 35. Site detection, annotation and event extraction from individual DCs in Exchange PAINT data*** *The workflow of the part of the Exchange PAINT processing pipeline used for extracting events from sites in individual DCs by determining the detected sites' positions in individual DCs, filtering probes based on geometrical constraints and finally extracting events in the different imaging channels originating from a given site using clustering (scale bars = 50nm).*

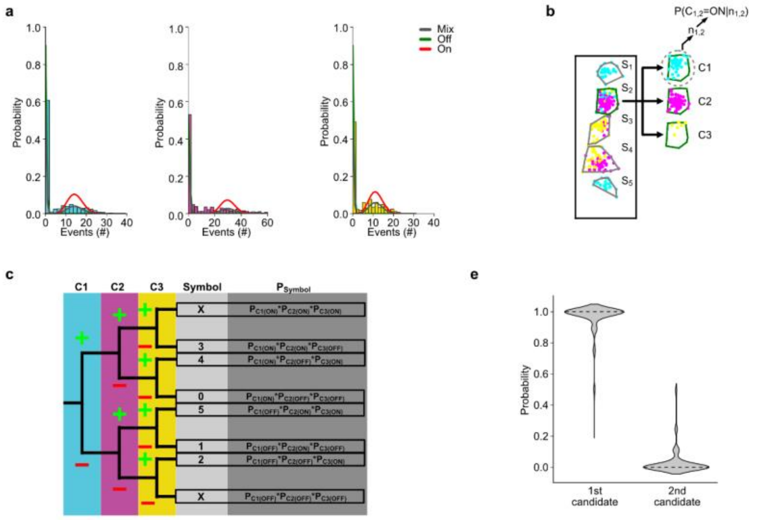


***Supplementary Fig 36. Site identity determination using a probabilistic approach a,*** *Plots showing the event distribution per single sites from the different channels (C1=cyan, C2=magenta, C3=yellow) with the Poisson-distributions corresponding to the "ON" state (red) and "OFF" state fitted to extract mean event frequencies and baseline probabilities corresponding to these states.* ***b,*** *Events extracted from individual sites of individual DPs were used to calculate the conditional probabilities of all possible channel states of all possible sites ($P(C_{i,j}=ON|n_{ij})$) using the mean rates and the observed event counts for the different channels ($n_{i,j}$).* ***c,*** *Decision tree showing the calculation of the conditional probabilities of individual sites on individual DCs that were calculated from the conditional probabilities of the channel states.* ***d,*** *Violin plot showing the distribution of calculated probabilities of DC codes for the highest probability candidates (1$^{st}$ candidate) and the second highest probability candidates (2$^{nd}$ candidate).*
